# The ratio of exhausted to resident infiltrating lymphocytes is prognostic for colorectal cancer patient outcome

**DOI:** 10.1101/2020.12.19.423379

**Authors:** Momeneh Foroutan, Ramyar Molania, Aline Pfefferle, Corina Behrenbruch, Axel Kallies, Terence P Speed, Joseph Cursons, Nicholas D Huntington

## Abstract

Immunotherapy success in colorectal cancer (CRC) is mainly limited to patients whose tumours exhibit high microsatellite instability (MSI). However, there is variability in treatment outcomes within this group, which is in part driven by the frequency and characteristics of tumour infiltrating immune cells. Indeed, the presence of specific infiltrating immune cell subsets has been shown to correlate with immunotherapy responses and is in many cases prognostic of treatment outcome. Tumour-infiltrating lymphocytes (TILs) can undergo distinct differentiation programs, acquiring features of tissue-residency or exhaustion, a process during which T cells upregulate inhibitory receptors such as PD-1 and loose functionality. While residency and exhaustion programs of CD8 T cells are relatively well-studied, these programs have only recently been appreciated in CD4 T cells and remain largely unknown in tumour-infiltrating natural killer (NK) cells. In this study, we use single cell RNA-seq data to identify signatures of residency and exhaustion in CRC infiltrating lymphocytes, including CD8, CD4 and NK cells. We then test these signatures in independent single cell data from tumour and normal tissue infiltrating immune cells. Further, we use versions of these signatures designed for bulk RNA-seq data to explore tumour intrinsic mutations associated with residency and exhaustion from TCGA data. Finally, using two independent transcriptomic data sets from patients with colon adenocarcinoma, we show that combinations of these signatures, in particular combinations of NK activity signatures, together with tumour-associated signatures, such as TGF-β signalling, are associated with distinct survival outcomes in colorectal cancer patients.

## Introduction

Immune checkpoint blockade (ICB) in colorectal cancer (CRC) has shown clinical benefits in a subset of CRC patients with high microsatellite instability (MSI) or deficient mismatch repair (dMMR), and in 10% of patients with microsatellite stable (MSS) or proficient MMR (pMMR) [1]. However, due the complex interplay between tumour-immune cells in the context of cancer immunotherapy, MSI status alone is often not enough to precisely predict response to ICB [2]. The main factors impacting the outcome of immunotherapy, and in general, survival outcome in patients, are the frequency and the characteristics of the immune cells in the tumour microenvironment (TME) [3–5].

Recent single cell studies in several cancer types have revealed a high level of heterogeneity in tumour infiltrating lymphocytes [6–8]. Importantly, a subset of CD8 T cells undergo “exhaustion”, a process that has been reported in activated T cells in response to persistent antigen exposure in the TME, resulting in a loss of functionality and expression of co-inhibitory receptors such as PD-1 or CTLA-4 [9]. Heterogeneity within the exhausted T cell population ranges from progenitor or precursor cells that exhibit self-renewal capacity to terminally differentiated cells [10–14], and this heterogeneity has been shown to impact the success of immunotherapy [15–17].

T cell exhaustion has largely been studied in the context of melanoma and non-small cell lung cancer (NSCLC) [7, 18, 19]. However, studies looking at T cell differentiation states and their tumour reactivity have yielded conflicting outcomes. Li et al found high reactivity of exhausted T cells towards tumour cells (measured by IFNg and TNFa production *ex vivo)* and showed high proliferation of exhausted cells at earlier stages of their exhaustion program [19]. Huang and colleagues, presented evidence that a subset of circulating CD8 T cells which express markers of exhausted cells (e.g. *PDCD1, CTLA4,* and *HAVCR2)* are the most proliferative T cells following ICB therapy, and that high ratio of exhausted CD8 T cell reinvigoration to tumour burden is prognostic in head and neck cancer patients [20]. Others have reported a low ratio of exhausted T cells to early activated cells to be associated with better response to ICBs [16].

Perhaps the most comprehensive study of CRC infiltrating immune cells was performed by Zhang et al [8, 21]. They showed that a TH1-like CD4 T cell cluster (expressing *CXCL13* and *BHLHE40)* share expression with some of the exhaustion genes found in exhausted CD8 T cells (with high *LAYN* transcript abundance). Interestingly, flow cytometry and single cell analyses in several cancer types have identified the presence of CD8^+^ FOXP3^+^ cells, which was further corroborated by T cell receptor (TCR) lineage tracing, suggesting a conversion between CD8 exhausted T cells and FOXP3^+^ T cells [7, 8, 19, 22]. These factors highlight the complex dynamics amongst TILs. In general, the presence of tissue resident memory T cells (T_rm_) has been reported to be associated with beneficial survival outcome and better response to immunotherapy in several cancer types [23–26]. While several studies have explored residency programs (or molecular signatures associated with residency) in T cells, these processes remain poorly understood in NK cells. Tissue-resident NK cells in steady-state are clearly distinguishable from conventional NK cells and have been referred to as type 1 innate lymphoid cells (ILC1s) [27]. We have previously observed ILC1s in murine tumour models and suggested these were derived from infiltrating NK cells which differentiated into ILC1s under conditions of high TGF-β signalling in the TME, permitting tumour cell escape from immunosurveillance [28].

TGF-β signalling not only plays critical roles in differentiation, development, and maintenance of various resident immune cells [29–32], but is also one of the key drivers of epithelial-mesenchymal transition (EMT) in cancer cells [33], which could impact the interplay between tumour cells and infiltrating immune cells. Active TGF-β cytokine is required for the differentiation of CD8 T cells into T_rm_ cells in several tissues [30, 34, 35]. Some studies have shown that TGF-β signalling is associated with T cell exclusion and tumour escape from immunosurveillance. Specifically, TGF-β signalling in fibroblasts of metastatic urothelial cancer patients is associated with T cell exclusion and reduced response to therapy with ICBs. In these patients, TGF-β blockade increased T cell infiltration and resulted in reduced tumour progression [36]. Similarly, another study in mouse models of CRC showed better ICB response after blockade of TGF-β signalling, driven by increasing T cell infiltration of the tumours and promotion of a T_H_1 phenotype [37]. Due to the crucial roles of TGF-β signalling in differentiation of resident cells, T cell exclusion in tumours, and its associations with survival outcome and response to ICBs, it is of utmost importance to better understand the relationship and interplay between TGF-β and immune or tumour cells.

This study, for the first time, attempts to tease apart transcriptional programs underpinning lymphocyte residency and exhaustion tumor-infiltrating T and NK cells using integrated analyses of single cell data from CRC and normal gut, as well as bulk transcriptomics data sets. Furthermore, we identify numerous tumour intrinsic mutations associated with these programs, in particular gene mutations associated with the NK exhaustion program, with several genes involved in cancer immune evasion. We then use our residency and exhaustion signatures in combination with other tumour associated programs (e.g. TGF-β signalling) to identify patients who have better survival outcome or may benefit from ICB immunotherapies. Specifically, we identify improved survival outcome for patients with high exhaustion and low residency programs in NK cells, and patients with high exhaustion programs in both CD8 and NK cells and low TGF-β signalling. This work has important implications for cancer immunotherapy as it suggests that strategies to prevent tumor residency may improve NK cell and CD8 T cell tumor immunity and patient outcomes. Collectively, we propose a new model that links distinct transcriptional programs to cancer patient survival and establish a new method that serves as valuable tool for researchers and health care professionals in analysing genomic and transcriptomic tumour biopsy data and interpreting tumor immune responses.

## Results

### Residency and exhaustion signatures define developmental trajectories of infiltrating immune cells

To establish residency and exhaustion signatures, we made use of Smart-seq2 data from tumor-infiltrating CD8, CD4 and NK cells generated by Zhang et al [21]. We first defined initial residency and exhaustion signatures by calculating the correlation between selected canonical residency or exhaustion markers and approximately 1300 genes collected from the literature, which were associated with these functional programs (see Methods; Supp. Figure S1 summarises the workflow for this study). We then used these initial signatures (Supp. Table S1) with singscore [38] to quantify the relative concordance of single-cells against these signatures, where a high score represents higher relative expression of genes within a given signature (Figure 1A). While residency scores for each cell type clearly separated immune cells in peripheral blood from those infiltrating tissue (either tumour or adjacent normal samples; across the x axis in Figure 1A), the exhaustion scores further separated cells with differences in their relative expression of exhaustion genes (across the y axis in Figure 1A). We then obtained genes that were differentially expressed between exhausted cells (with high exhaustion and low residency scores) and resident cells (with high residency and low exhaustion scores; *thresholds shown in Figure 1A),* and further filtered the resulting genes by group comparisons with percentile thresholds (see Methods).

**Figure 1.**
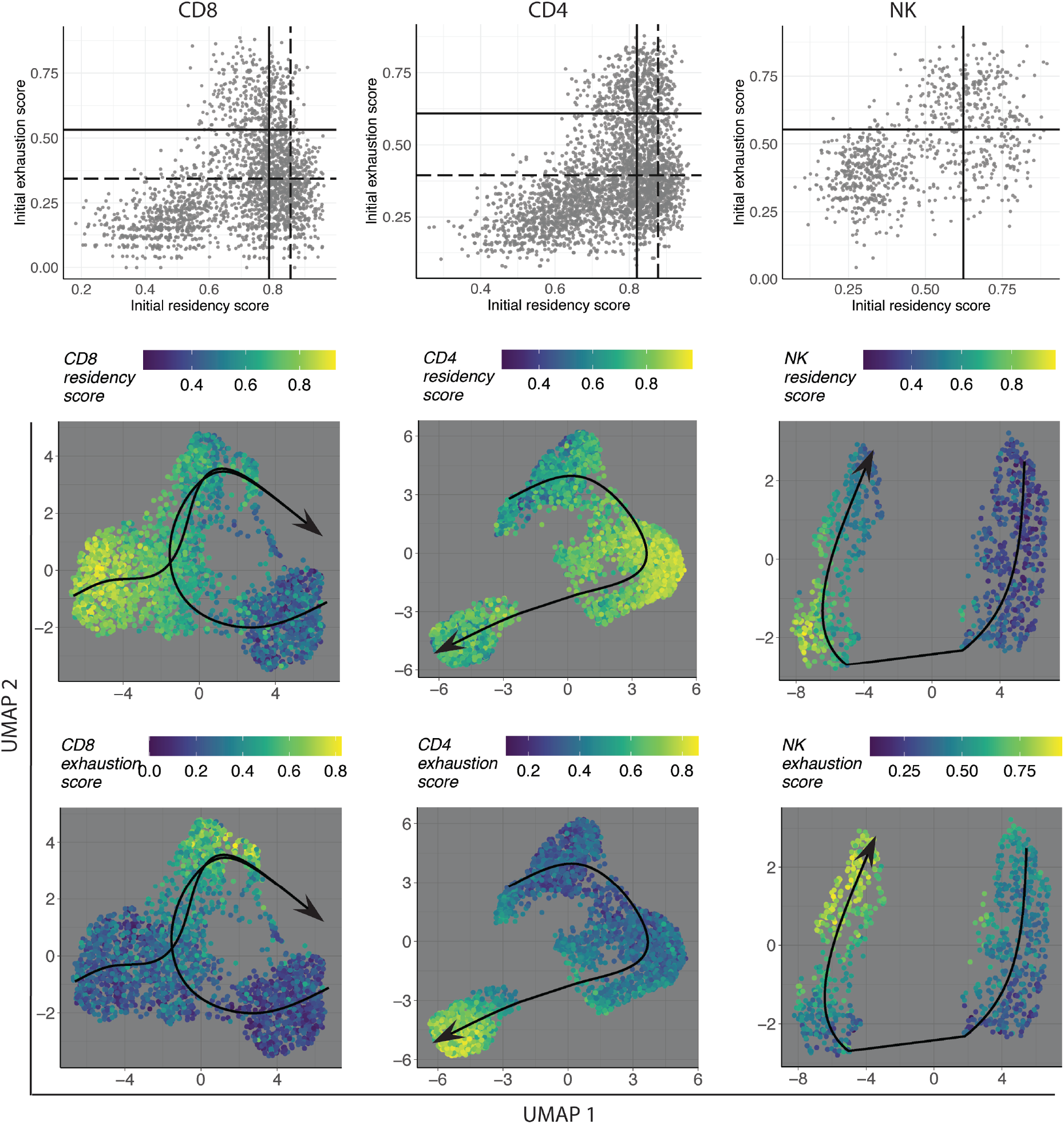
(Top row) Initial exhaustion and residency scores in each of the CD8, CD4, and NK subsets of the Zhang Smart-seq2 data. Solid lines show the thresholds used to define Exh cells in CD8 and CD4 cells, and dashed lines represent the thresholds used to define Res cells. (Bottom panel) Clustering and UMAP performed on our marker genes in each of the cell populations, coloured by Exh and Res signature scores. Lines represent the trajectory analysis performed by slingshot.

The resultant gene sets were residency (Res) and exhaustion (Exh) signatures for each of the CD8, CD4 and NK cells that can be used for *infiltrating immune cells* (called CD8_Res *[N_Genes_* = 43], CD8_Exh *[N_Genes_* = 27], CD4_Res *[N_Genes_* = 31], CD4_Exh *[N_Genes_* = 83], NK_Res *[N_Genes_* = 61] and NK_Exh *[N_Genes_* = 19]; Supp. Table S2). We further merged all the Res signatures together to define a general Res signature (All_Res; *N_Genes_* = 100) and all the Exh signatures together to define a general Exh signature (All_Exh; *N_Genes_* = 118). The single cell scores from these signatures could identify a subset of cells with higher evidence of Exh or Res programs in the Zhang Smart-seq2 data (Supp. Figure S2), with exhausted NK cells largely restricted to tumours with MSI status. In general, our results showed that Exh programs were similar between CD8 T cells and NK cells, however, Res signatures were more similar between CD8 and CD4 cells. Zhang et al reported more similarity between T_H_1 like clusters of CD4 cells with exhausted CD8 cells [8] but in our analysis, we observed higher exhaustion scores in a large number of tumour specific CD4 Tregs, which is more similar to the CD4 dysfunctional Treg signature in melanoma samples reported by Li et al [19].

While *HAVCR2* (encoding TIM3) was the only gene shared across the Exh signatures for all three cell types, we identified ten overlapping genes across all the Res signatures (Supp. Table S2). A small number of genes also overlapped between each pair of signatures. Some of the transcription factors (TFs) that we identified in the All_Res signature included genes associated with activating protein 1 (AP-1) TFs, which are downstream of TCR activation, NF-κB signalling, and early growth response downstream of AP-1 and NF-κB signalling (Supp. Table S2). Although it was identified as one of the CD8 Res markers, *GPR15* is one of the less studied GPCRs that also showed restricted expression in CD4 and NK Res cells (Supp. Figure S3). From the full list of genes in the All_Exh *(N_Genes_* = 118) signature, we identified three genes associated with G-Protein coupled receptors (GPCR) signalling (e.g. CCR8) and ten transcription factors (Supp. Table S2).

Consistent with the idea that exhaustion signatures show a strong degree of overlap with activation signatures, a small subset of the Exh cells from all the three cell types exhibited a high level of proliferation. Using a proliferation signature and the Exh signatures obtained here, we further identified signatures for proliferative Exh cells (see Supp. Figure S4 and Supp. Table S3 for the list of DEGs).

After clustering and dimensionality reduction on our cell type specific marker genes, we performed unsupervised trajectory analyses, which could not only separate blood from tissue infiltrating immune cells, but also showed resident cells as an intermediate step between naïve cells (mostly in blood circulation) and exhausted cells (Figure 1B). Interestingly, while the Exh subset of CD4 and NK cells are a down-stream lineage of Res cells in tissues, the Exh subset in CD8 T cells seem to derive from both tissue-infiltrating Res cells as well as circulating CD8 T cells.

To further examine the development of NK cells in tumor tissue, we assessed the expression of the several genes defining NK cell maturation and tissue residency [39] in the NK cells extracted from tissue and blood (Supp. Figure S5). Interestingly, we observed a population of tissue-infiltrating NK cells (annotated as the GZMK population by Zhang et al), with expression of naïve markers *(CCR7, SELL,* and *GPR138)* as well as inhibitory markers (at least one of *PDCD1, TIGIT, HAVCR2,* or *CTLA4;* Supp. Figure S6). Overlaying expression of these genes on the trajectory plots identified these cells as belonging to an intermediate stage of the trajectory (Supp. Figure S6).

To examine the association between these programs and TGF-β signalling, we also scored single-cells against a TGF-β signature for tissue-resident memory T (T_rm_) cells reported by Nath et al [32]. Our results showed higher correlations between TGF-β signature scores and Res scores in CD8 and NK cells compared to Exh scores, while we observed higher correlation between the CD4_Exh and TGF-β program (Supp. Figure S7). Most of the CD4_Exh cells which also have high TGF-β scores are tumour associated Tregs, and TGF-β is important for the Treg infiltration, retention, and suppressive function [40].

### Transcriptional signatures identify residency and exhaustion in single cell data from CRC and healthy tissue resident cells

Using the signature genes obtained above, we generated UMAP plots in three independent single cell data sets, including two CRC data sets and one normal tissue data set: 10X data from Zhang et al [21], the single-cell data from de Vries et al [41], as well as a normal human gut data from James et al [42] (See Method section; Figure 2A). Similar to the Zhang Smart-seq UMAPs, the first dimension of the UMAP plots in the Zhang 10X data generated using our marker genes separated the immune cells in peripheral blood from those in the tissue. We showed that while a subset of cells only exhibited evidence for Exh or Res, some cells displayed a ‘hybrid’ program. In normal human gut we observed relatively more Res cells, which confirms previous observations that showed more Res cells in normal samples [8]. Interestingly, we also observed a small subset of exhausted cells in normal gut tissue, perhaps due to the persistent presence of microbiota-derived antigens in the gut.

**Figure 2.**
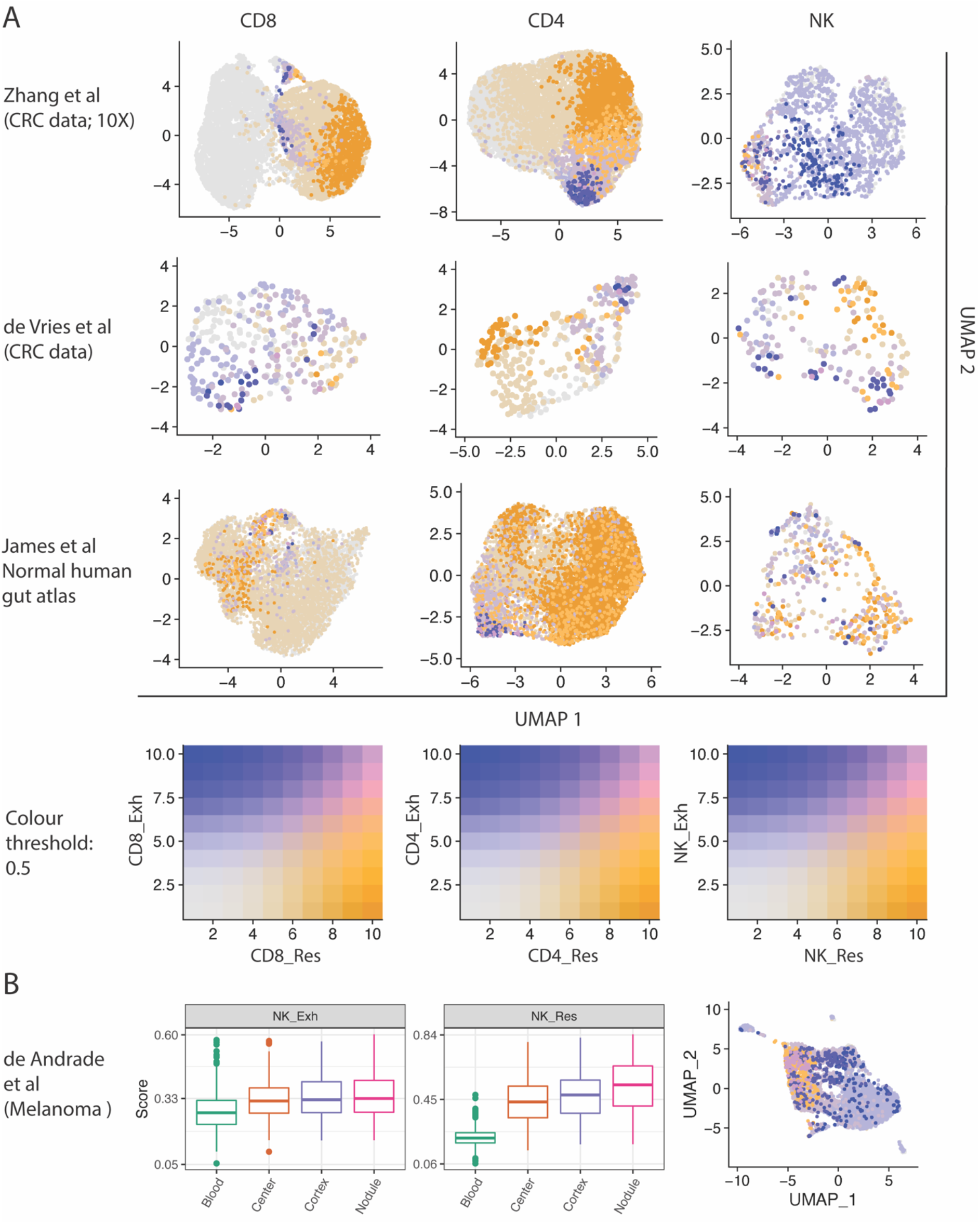
Test signatures in independent single cell data sets of cancer and normal samples. (A) UMAP blend plots of CD8, CD4, and NK cells generated using signature genes in two CRC single cell data (top and middle row), and normal gut cells (bottom row). Scores were scaled for better visualisation of both scores on the same plot. (B) Boxplots and UMAP blend plot for the NK cells extracted from a melanoma patient, coloured by the scaled NK_Exh and NK_Res scores.

As the frequency of NK cells was low in some of the test data sets, we further examined our NK markers in ~4,000 NK cells extracted from a melanoma patient [43] (Figure 2B). Using this data set, we found that NK cells infiltrating in a distinct tumour nodule (in the cortex of the tumour) had the highest NK_Res score, followed by NK cells within the cortex (margin) and centre (core) of the tumour. NK cells from peripheral blood exhibited the lowest NK_Res score as expected. Although NK_Exh scores were also relatively higher in the NK cells extracted from the nodule compared to the NK cells from blood, the differences between NK_Exh scores from different sources were not as high as those in NK_Res scores. This confirms the observed evidence of NK_Exh across NK cells extracted from different sources in the CRC single cell test data sets (Figure 2A & B), and suggests that NK cells from different sources (blood, cortex, centre, and nodule) in cancer patients may exhibit features of exhaustion, similar as it has been reported recently in patients with chronic hepatitis B [44].

### Exhaustion and residency programs are enriched in MSI/dMMR samples and CMS1/CMS4 subtypes

As tumour samples from patients include both cancer cells and non-cancer cells from the tumour microenvironment (such as infiltrating immune cells), we further refined our signatures to only retain genes with low expression in tumour cells. This increases our confidence when scoring bulk tumour samples, ensuring that scores are primarily defined by infiltrating immune cells and not tumour cells. To this end, we examined the expression of all of our marker genes in the CRC cell lines from the Cancer Cell Line Encyclopedia (CCLE) and CRC laser capture microdissected (LCM) cells from Tsukamoto et al (GSE21510) [45], and retained genes that showed low expression in either of the two data sets (see Method section). Therefore, these signatures can be used for separate immune cells from *tumour samples* and are named with “Bulk” suffix (e.g. CD8_Res_Bulk; Figure 3 and Supp. Table S2).

**Figure 3.**
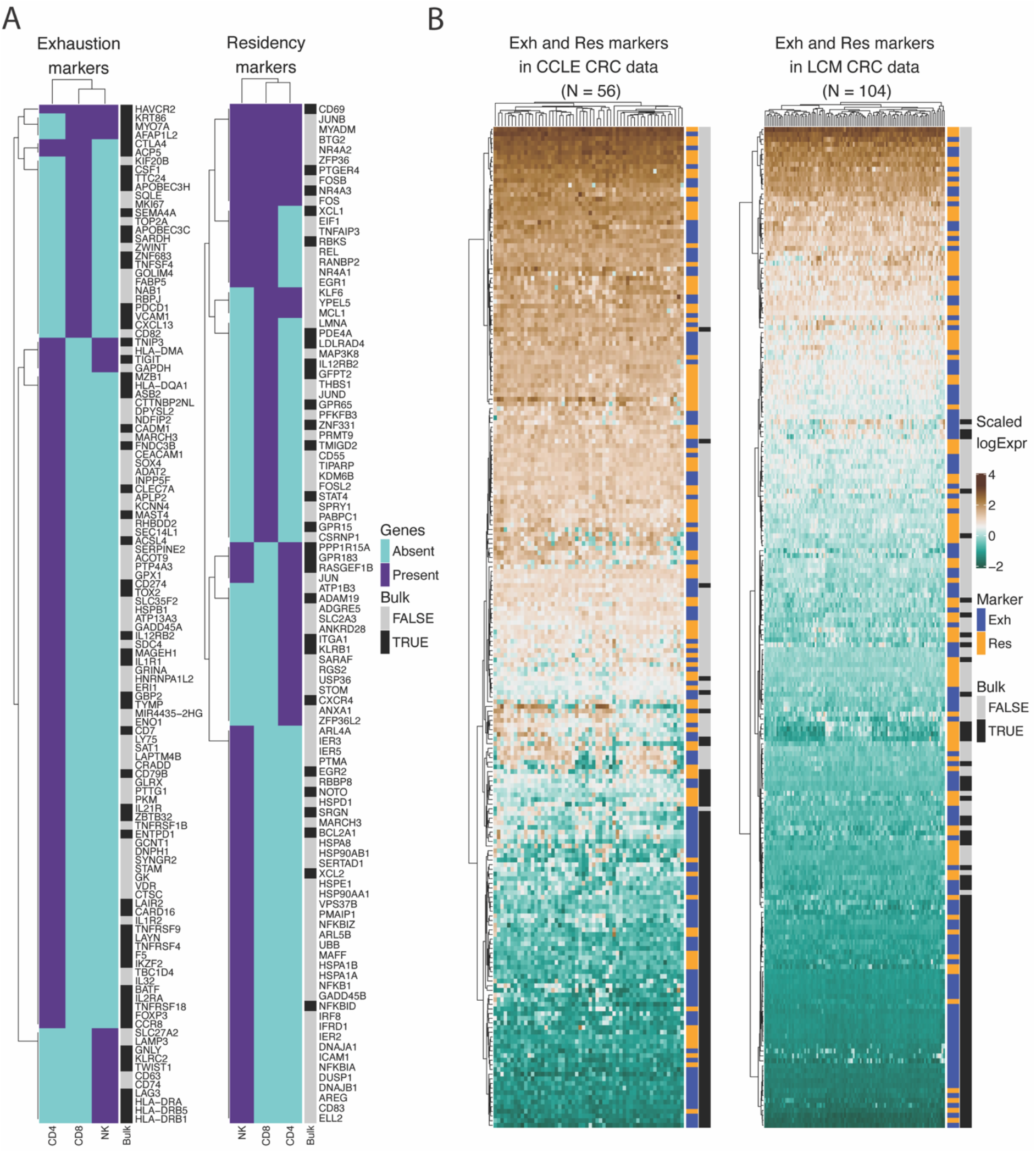
Exhaustion and residency markers and their refinement in two cancer data sets CCLE and LCM. **A)** Hierarchical clustering of marker genes from different cell types; genes are annotated based on whether or not they pass the “bulk tumour threshold” in CCLE and LCM expression data. **B)** Expression of Res and Exh markers (coloured orange and blue) in the CCLE and LCM data; genes annotated in black are those that pass the bulk tumour thresholds in at least one of the two data sets.

We then used these signatures in colon cancer patient samples from two independent datasets: RNA-seq from TCGA (*N_sampies_* = 479, including 40 matched normal samples), and microarray from Marisa et al [46] (*N_samples_* = 585, including 19 matched normal samples). Stratifying samples based on scores and MSI/dMMR status showed relatively higher exhaustion and residency scores in MSI-H and dMMR samples, although the difference between MSI-H/dMMR and MSS/pMMR was relatively larger for Exh scores compared to Res scores (Figure 4A). We then divided samples based on the four consensus molecular subtypes (CMS), and in both tumour datasets we observed relatively higher Exh and Res scores in CMS1 and CMS4 subtypes (Figure 4B), which is consistent with the characteristic of these subtypes (Higher number of MSI samples and higher immune activation in CMS1, CSM4 and CMS2/3, respectively) [47, 48]. As the CMS1 subtype has higher occurrence in proximal CRC samples (right-side), we further confirmed higher scores, specifically for NK_Exh_Bulk signature, for proximal (right) compared to distal (left) tumours (Supp. Figure S7).

**Figure 4.**
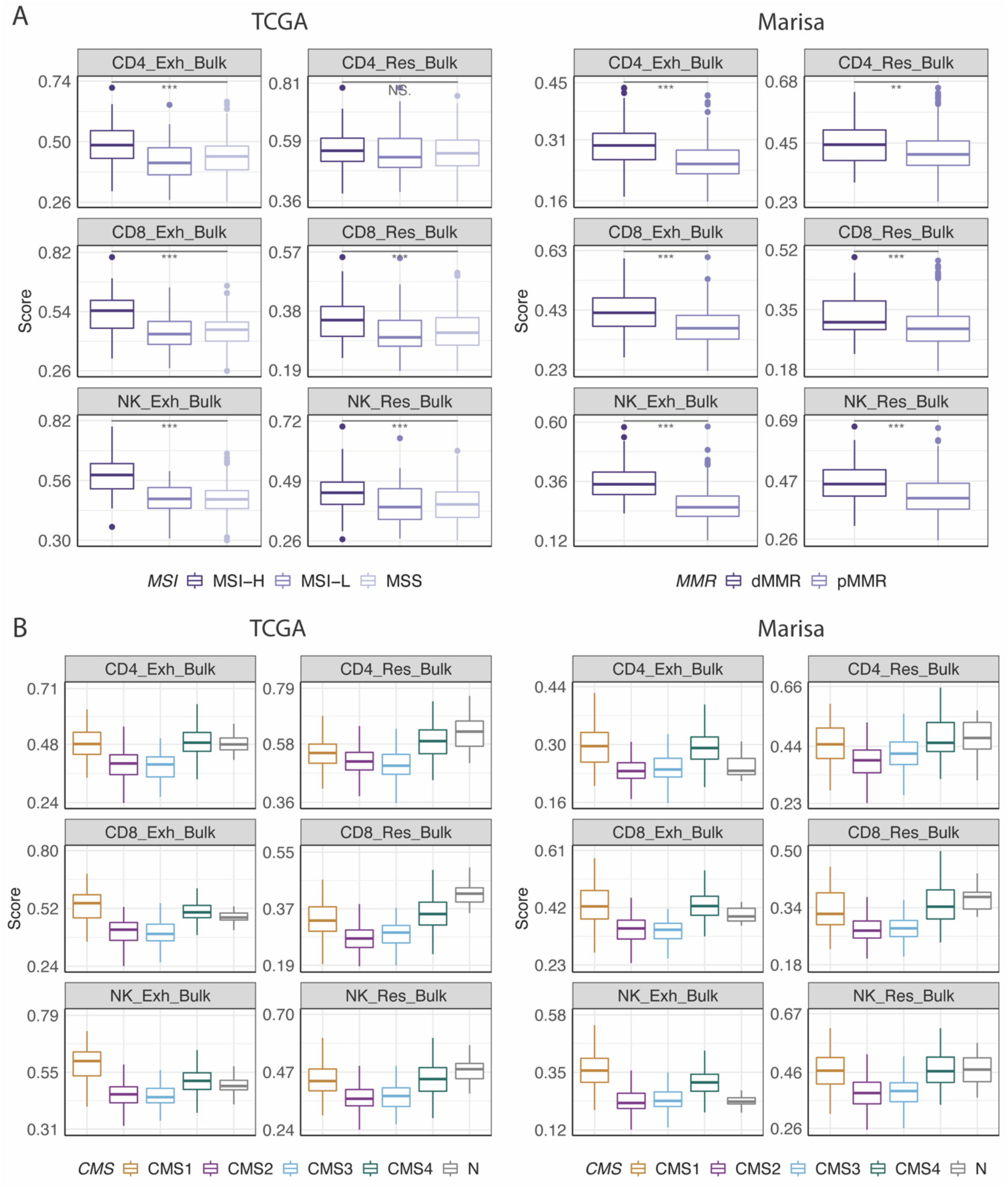
Exh and Res bulk signature scores across the TCGA and Marisa samples stratified by their MSI or MMR status (**A**) and by their CMS subtypes (**B**). MSI samples, as well as CMS1 and CMS4 subtypes were associated with higher Exh and Res scores.

Consistent with higher Res scores in single cell data from normal gut samples (Figure 2), we also observed relatively higher evidence of the Res program in normal bulk samples (scores were equivalent or higher than the scores in CMS1 and CMS4), with relatively lower Exh scores, specifically for CD8 and NK cells. These results not only are consistent with previous research that showed higher immune infiltration in MSI-H and CMS1 samples [47, 48], they also suggest that the presence of exhausted cells is more cancer specific, with relatively less enrichment in normal tissues.

### Tumour intrinsic mutations associated with exhaustion and residency

In order to identify tumour intrinsic mutations associated with residency and exhaustion programs, we used somatic mutations data from the TCGA COAD, and performed two types of analyses. In the first analysis, we generated linear models, examining the association between each gene mutation and each Exh_Bulk and Res_Bulk score. In general, we found more gene mutations associated with exhaustion signatures than with residency programs (Supp. Table S4), with the highest number related to NK_Exh_Bulk score (*N_Genes_* = 3371),followed by CD8_Exh_Bulk (*N_Genes_* = 2130). Among the residency signatures, CD8_Res_Bulk scores were associated with the highest number of gene mutations (*N_Genes_* = 928), followed by NK_Res_Bulk score (*N_Genes_* = 832).

Amongst the list of significant genes, we found several well-known CRC gene mutations. Mutant BRAF showed significantly higher scores for all Exh_Bulk and Res_Bulk signatures, except for CD4_Res_Bulk. APC mutations, on the other hand, were associated with lower Exh_Bulk and Res_Bulk scores, with the difference being more profound for Exh_Bulk scores compared to Res_Bulk scores (Figure 5). These results are consistent with previous research that showed relatively more MSI samples with *BRAF* mutations and more *APC* mutations in MSS samples [48, 49]. BRAF not only is associated with MLH1 methylation (responsible for DNA mismatch repair) and developing MSI samples [50], but mutations in this gene (specifically V600E) can also impact tumour hypoxia across a number of cancer types [51–53].

**Figure 5.**
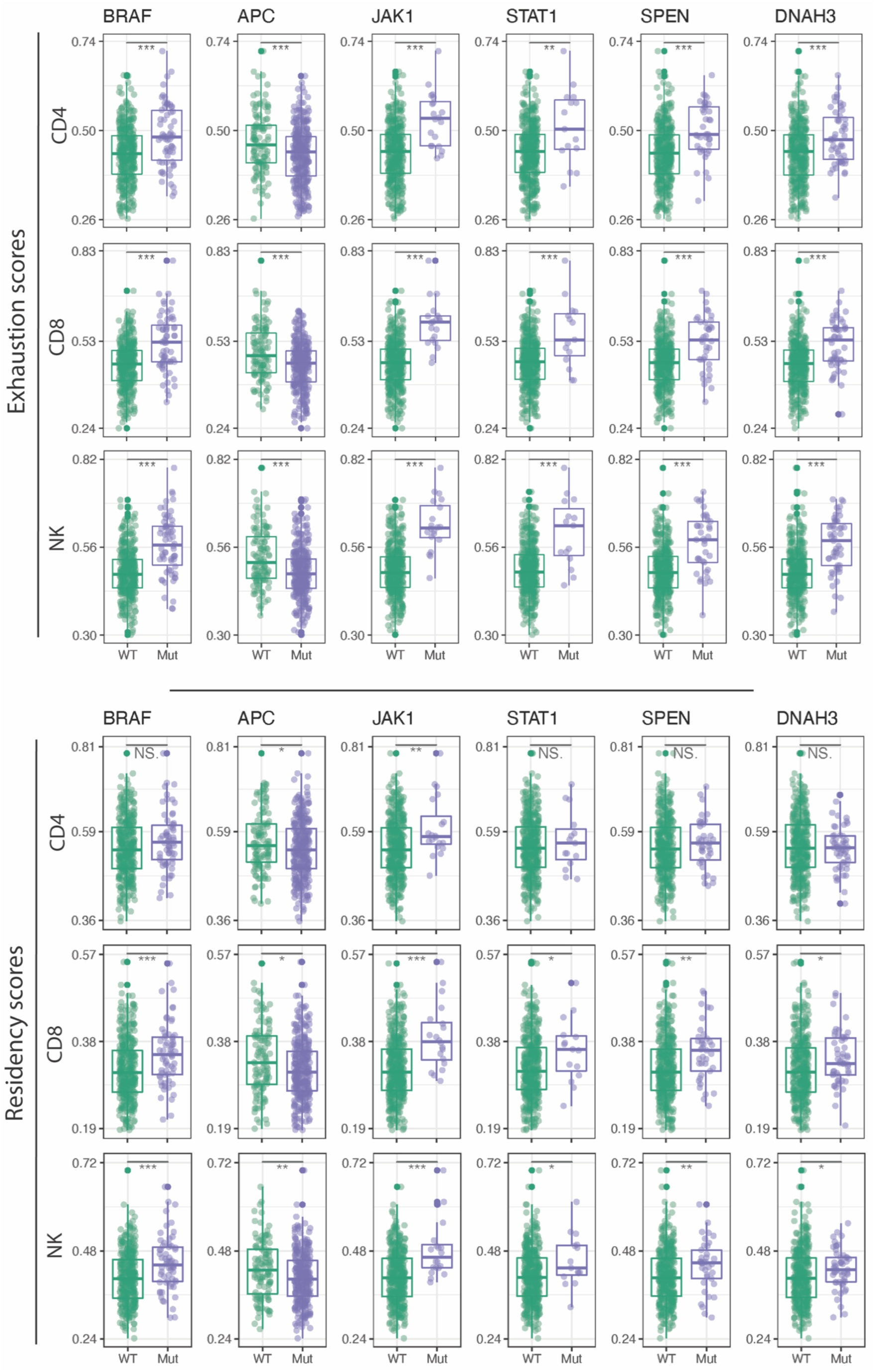
Exh_Bulk and Res_Bulk scores for each cell types for TCGA samples with mutant and WT forms of selected genes.

We also identified mutations in several genes impacting on tumor antigen presentation pathway JAK/STAT; while mutations in JAK1 were associated with both higher Exh_Bulk and Res_Bulk scores, STAT1 mutations had significantly higher Exh_Bulk scores but not Res_Bulk scores. Finally, our results identify novel genes associated with these programs, for example, SPEN (a transcriptional repressor, which synergizes with RUNX2) and several dynein axonemal heavy chain (DNAH) genes (components of microtubules, involved in ATP-dependent cell motility; Supp. Table S4). Focusing on top gene mutations (adjusted p-value < 0.0001) associated with the NK_Exh_Bulk program (*N_Genes_* = 1027), we identified several GO terms, such as actin filament-based processes, calcium ion transmembrane transport, regulation of GTPase activity, response to oxygen levels, response to TGF-β, and stem cell proliferation (Supp. Table S5 and Supp. Figure S9).

In the second analysis, we focused on NK cell signatures, which showed relatively higher gene mutations associated with exhaustion. We performed elastic net regression by including all the significant genes obtained in the first analysis (for each of the NK_Exh_Bulk and NK_Res_Bulk), as well as clinical factors (see Method section). This analysis resulted in 44 genes predictive of NK_Exh_Bulk; examples of these predictive genes are those associated with antigen presentations *(HLA-B, JAK1, TRRAP), POLD1,* and TGM6, which is less appreciated in this context. Mutations in all genes, except for the APC, were associated with higher NK_Exh_Bulk scores. Additionally, the CMS3 subtype showed a significantly negative coefficient for NK_Exh_Bulk, which further confirms that this subtype is an “immune excluded” subtype [48]. CMS4 was one of the predictive factors for both NK_Exh_Bulk and NK_Res_Bulk signatures. Indeed, CMS4 was the most predictive factor for NK_Res_Bulk scores (compared to all the genes and clinical factors included as predictors), and this is of interest as CMS4 is associated with EMT programs and TGF-β signalling, the latter playing an important role in differentiation of tumour infiltrating resident cells [29, 30]. We found 75 genes predictive of NK_Res_Bulk (mutations in 55 genes were associated with higher scores), of which 12 genes overlapped between the two signatures, including BRAF, APC, and HTR5A (Supp. Table S6, Supp. Figures S10–12). Other predictive genes of NK_Res_Bulk associated with TGF-β signalling and the mesenchymal-epithelial transition (MET) were *TGFBRAP1* and *HGF,* the latter is known to modulate immune response [54, 55]. Interestingly, we found *PAK4* as one of the predictive genes for NK_Res_Bulk. *PAK4* was recently introduced as a target in cancer immune evasion as its expression was inversely correlated with CD8 T cell infiltration and ICB response in melanoma, and *PAK4* overexpression correlated with an activated Wnt-β-catenin pathway [56, 57].

### Combined Exh and Res scores identify patients with distinct survival outcome

As heterogeneity amongst the infiltrating immune cells has been shown to be associated with response to immunotherapy and patient’s survival outcome, we examined whether the Res_Bulk and Exh_Bulk signatures have associations with survival outcome in patients. We used the overall survival (OS) data available in both TCGA and Marisa, as well as progression free interval (PFI) in TCGA and relapse free survival (RFS) in Marisa data [46]. We also note that these two data sets differ in their clinical definitions of survival outcomes; for example, TCGA data defines PFI based on the date of diagnosis, whereas Marisa et al define RFS based on the date of surgery. However, we analysed each of these data sets separately, and look for any consistency between the two sets of results. In addition to all the CD8, CD4, NK, and All Exh_Bulk and Res_Bulk signatures, we merged CD8 and NK Res_Bulk signatures (called CD8NK_Res_Bulk; *N_Genes_* = 23), and CD8 and NK Exh_Bulk signatures (called CD8NK_Exh_Bulk; *N_Genes_* = 26) to examine the role of the combined signatures of CD8 T cells and NK cells.

Focusing on OS, we first examined whether each individual Res or Exh score was associated with OS by dividing the samples into two groups with high and low scores based on median values and fitting Cox multivariate models to these groups. We identified better OS for samples with higher CD8NK_Exh_Bulk and NK_Exh_Bulk scores compared to those with relatively lower scores for these signatures in Marisa data (Supp. Figure S13A); however, we did not see significant associations for these two signatures in the TCGA OS data. We then stratified samples based on score pairs using 2-dimensional score landscape plots (Figure 6B), including Exh_Bulk and Res_Bulk signatures, as well as a number of tumours associated signatures (see Methods). The multivariate Cox analysis identified poorer OS for patients with low CD8NK_Exh_Bulk & high TGF-β signalling in both data sets (Figure 6C). Other consistent findings across the two datasets included poorer OS for patients with low NK_Exh_Bulk & high IL6_JAK_STAT3 signalling, and for patients with low NK_Exh_Bulk & low DDR (Figure 6A).

**Figure 6.**
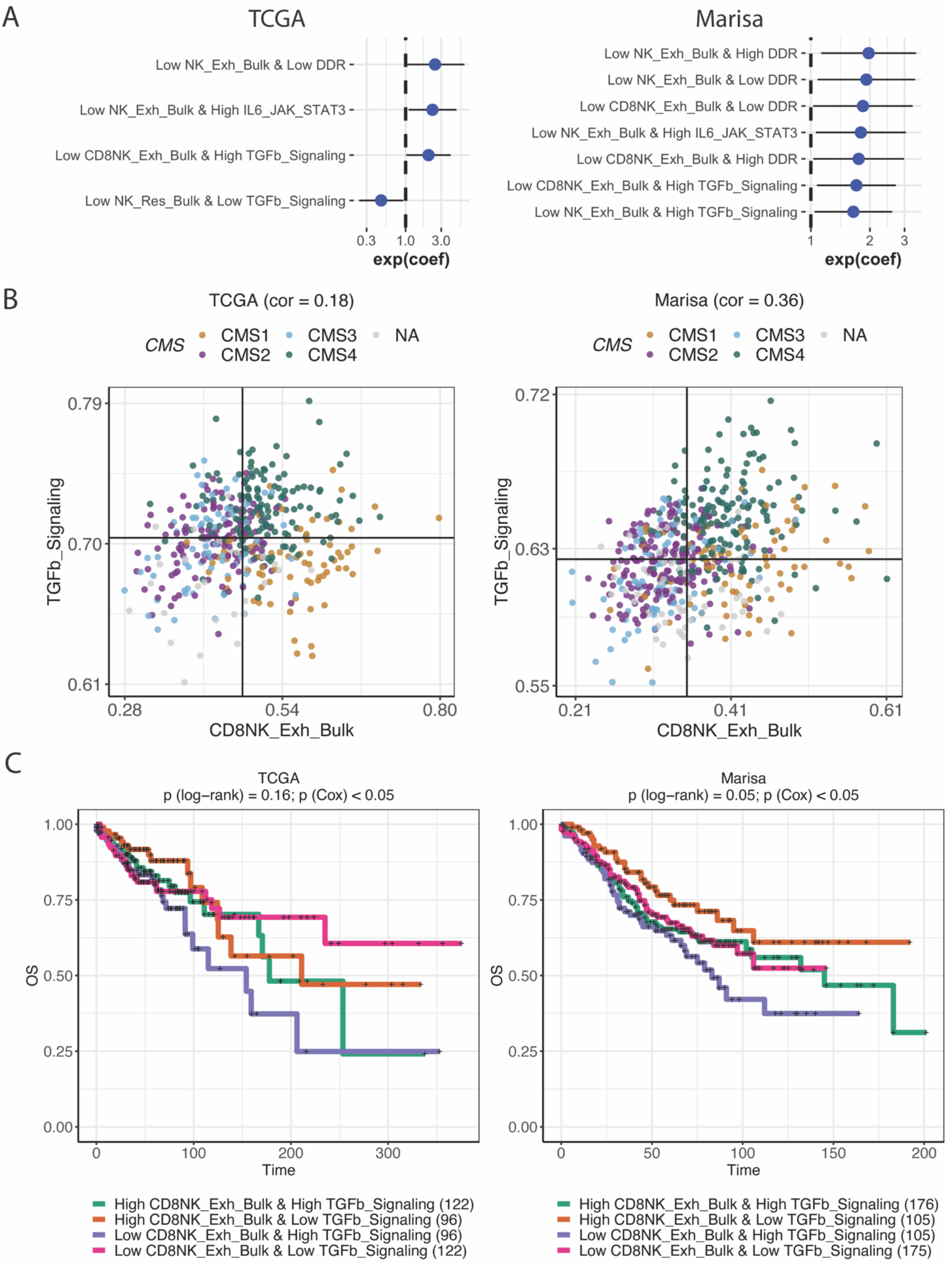
(**A**): Hazard ratios obtained from Cox multivariate models using score pairs (plus covariates) in the TCGA and Marisa OS data. (**B**) Score landscapes comparing CD8NK_Exh_Bulk scores vs TGF-β signalling scores in the TCGA and Marisa data, coloured by CMS subtypes. (**C**) Kaplan-Meier curves demonstrating overall survival (OS) for patients stratified based on scores shown in B.

We then compared the progression free interval (PFI) from TCGA and relapse free survival (RFS) from the Marisa data. Considering individual scores in multivariate Cox models, we identified a subset of patients that scored lower for the NK_Res_Bulk signature and were associated with better PFI in TCGA data (Supp. Figure S13B), although we did not find significant associations between NK_Res_Bulk scores and RFS in Marisa data. Stratifying samples based on score pairs and performing Cox models, we identified High CD8NK_Exh_Bulk & Low_NK_Res scores associated with longer PFI in TCGA, and consistent with this, samples with low CD8NK_Exh_Bulk & High NK_Res_Bulk scores showed lower RFS in Marisa data (Figure 7). We also found significant associations with RFS/PFI when combining our Res_Bulk and Exh_Bulk signatures with a number of tumour-associated signatures, such as DDR, TP53 signalling, and those involved in EMT program or metabolic processes (e.g. oxidative phosphorylation (OXPHOS) and peroxisomes; Figure 7A). In general, we observed better PFI/RFS for lower NK_Res_Bulk program when combined with lower scores for TP53 signalling, hypoxia/EGF-driven EMT, epithelial, or peroxisome scores. We also found poorer RFS/PFI for samples with high CD8NK_Res_Bulk and low TP53_neg scores, or samples with low CD8NK_Res_Bulk scores that also had lower DDR scores (Figure 7).

**Figure 7.**
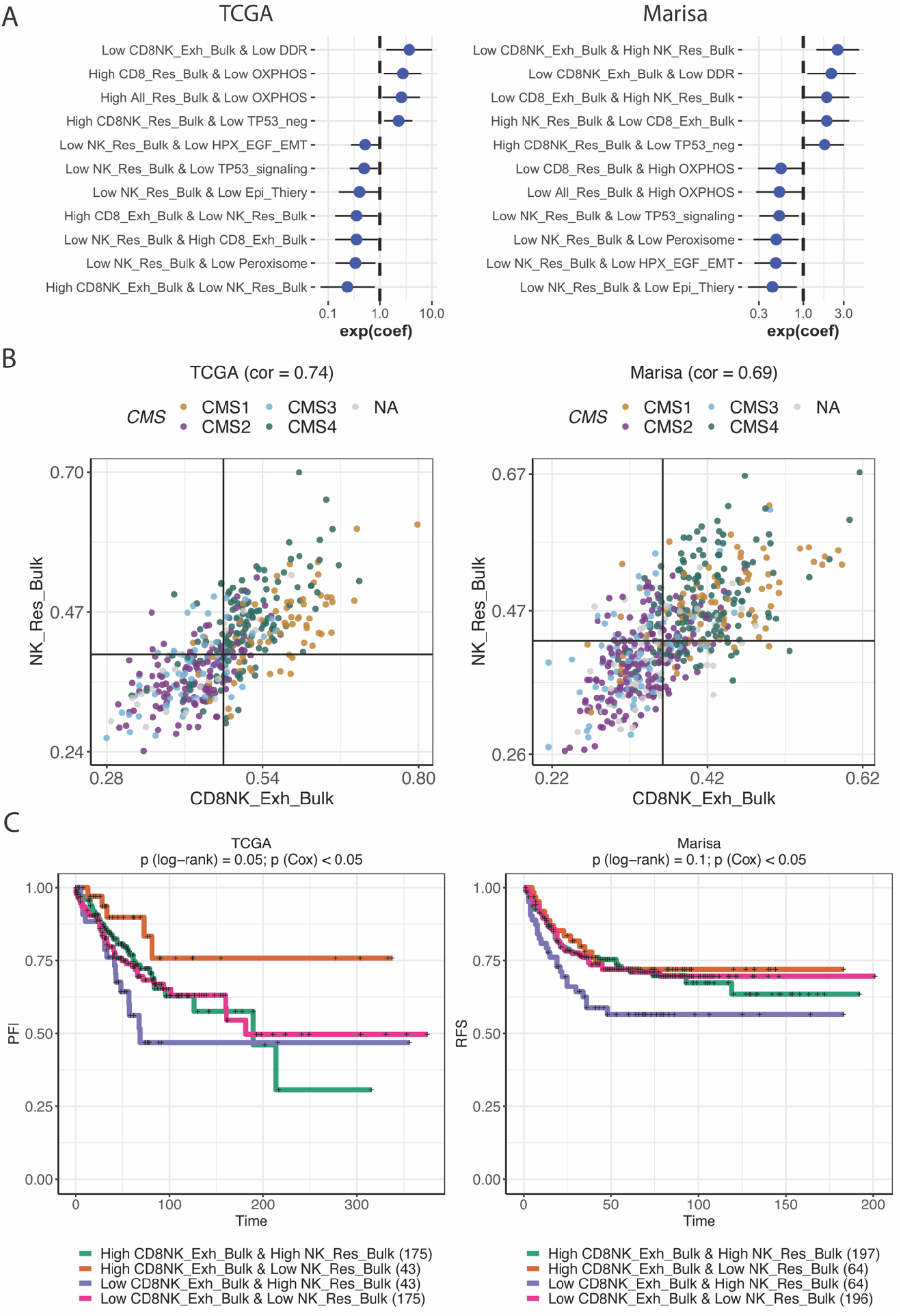
(**A**): Hazard ratios obtained from Cox multivariate models using score (plus covariates) in the TCGA PFI and Marisa RFS data. (**B**) Score landscapes comparing CD8NK_Exh_Bulk scores vs NK_Res_Bulk scores in the TCGA and Marisa data, coloured by CMS subtypes. (**C**) Kaplan-Meier curves demonstrating PFI or RFS for patients stratified based on scores shown in B.

While Res_Bulk and Exh_Bulk scores consistently showed positive correlations with TGF-β signaling, EMT (driven by TGF-β or hypoxia and EGF) and TP53-negative scores in both data sets, negative correlations were observed between Res_Bulk and Exh_Bulk scores and DDR, OXPHOS and TP53-positive scores (Supp. Figure S13C-D). However, when focusing on each of the cancer related scores, there were differences in their correlation with Res_Bulk and Exh_Bulk scores. For example, consistent with our findings in single cell data, we observed higher correlation between TGF-β signalling scores and Res_Bulk scores compared to those from TGF-β signalling scores and Exh_Bulk scores (Supp. Figure S13C-D).

For each of the OS and PFI/RFS data, we also aimed to find patients with more distinct programs of Exh_Bulk and Res_Bulk, by subsetting the data into patients with high Exh_Bulk and low Res_Bulk scores and those with low Exh_Bulk and high Res_Bulk scores, and performed the Cox multivariate analysis on these subsets. We found that, in both the TCGA and Marisa data, samples with low NK_Res_Bulk & high NK_Exh_Bulk showed improved OS and longer PFI/RFS time compared to those with high NK_Res_Bulk & low NK_Exh_Bulk scores (Figure 8). Similarly, samples with high CD8_Exh_Bulk & low NK_Res_Bulk showed better survival outcomes in both data sets (Supp. Figure S14). We noted that there was a relatively small subset of samples with opposite characteristics regarding Exh and Res programs, which is due to the positive correlations between the two signatures. Considering that there were no genes overlapping between the two signatures, this observation highlights the true biological associations between these two programmes. Interestingly, samples with opposite characteristics included a mix of different CMS subtypes (Figure 8A) and different MSI status (data not shown), suggesting that our scoring approach adds a new layer of information for sample stratification, and Exh and Res scores are predictors of survival outcome in the COAD patients, independent of known clinical variables.

**Figure 8.**
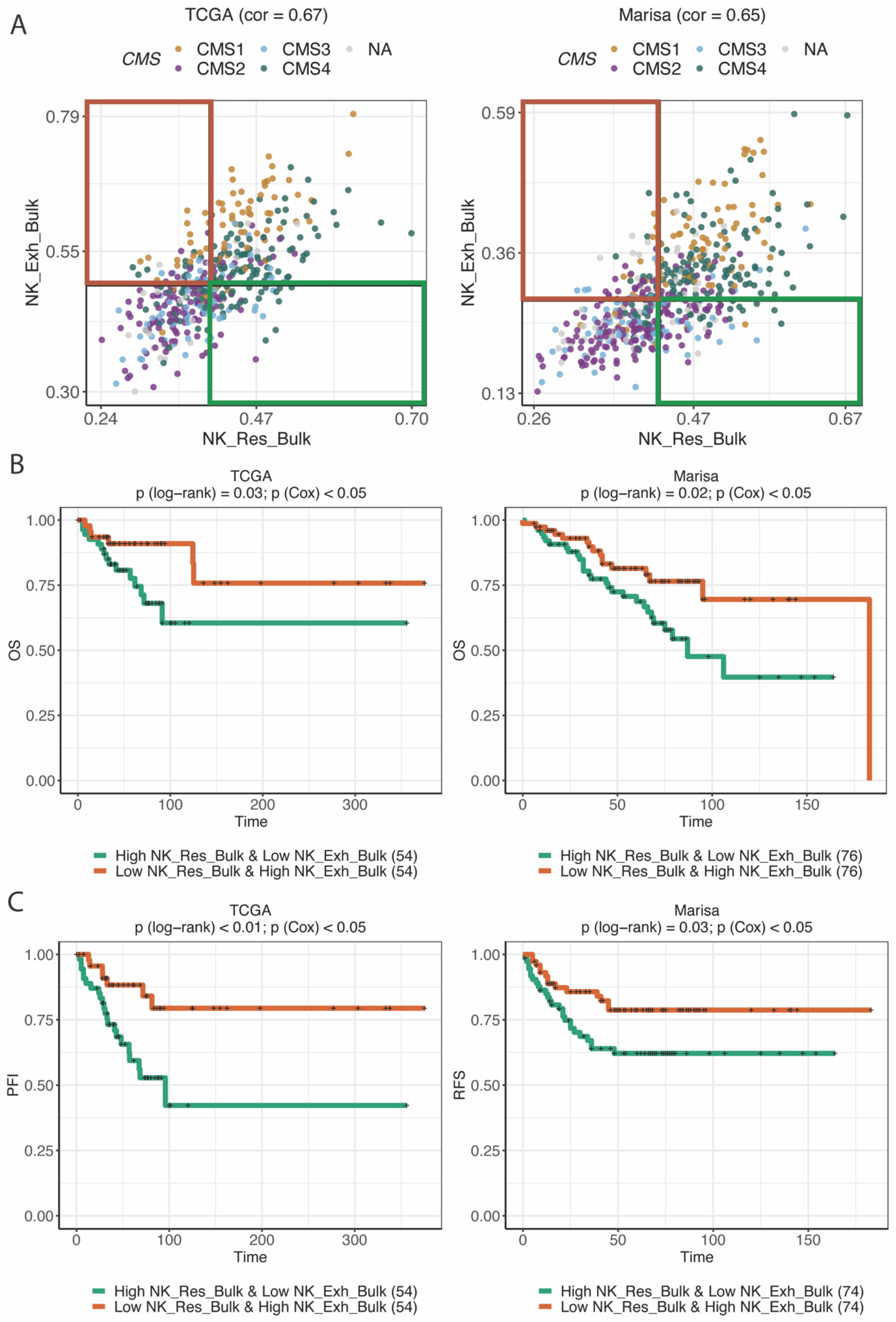
(**A**) Score landscapes comparing NK_Exh_Bulk scores vs NK_Res_Bulk scores in the TCGA and Marisa data, coloured by CMS subtypes. (**B**) Kaplan-Meier curves demonstrating OS for patients stratified based on scores shown in A (samples in coloured boxes); (**C**) Kaplan-Meier curves demonstrating PFI or RFS for patients stratified based on scores shown in A.

## Discussion

In this study, for the first time to our knowledge, we have leveraged single cell and bulk transcriptomic data to distinguish exhaustion and residency programs in both T cells and NK cells in CRC, validated them across several independent data sets, and shown associations with tumour genomic profiles and patient survival outcome.

Several genes in the Exh and Res signatures have been previously reported as part of the exhausted/dysfunctional or resident T cell signatures in different cancer types [7, 8, 18, 19], and some have received less attention in this context. We identified members of the APOBEC family of single-stranded DNA cytosine deaminases in our CD8_Exh signatures *(APOBEC3H and APOBEC3C),* which have been shown to share similar functions in the context of HIV and cancer [58]. Indeed, some APOBEC genes are expressed in tumour infiltrating T cells and are associated with better clinical outcome [59, 60]. GAPDH, one of our identified Exh markers, induces the translation of IFNG and IL2 in highly glycolytic T cells and suppresses translation of key genes in CD4 T cells following inhibition of glycolysis [61]. Additionally, GAPDH has been previously shown to be upregulated in rapidly cycling and activated NK cells compared to slowly cycling NK cells [62]. One of the CD4_Exh genes associated with GPCR was CCR8, and of note, combining anti-CCR8 with other immunotherapy agents in CRC has been shown to prevent the suppressive effect of CD4 Tregs and improve patients’ survival outcome [63]. GPR15, one of our CD8_Res markers, which also has restricted expression in Res CD4 and NK cells, is a GPCR involved in T cell trafficking in colon, and its expression in Tregs is promoted by TGF-β, mediating the suppression of inflammation in the colon [40]. CXCR4, identified as one of our CD4_Res genes involved in GPCR signalling, was just recently suggested to mediate immune suppression in pancreatic and CRC samples, and the use of anti-CXCR4 in MSS samples increased the immune response [64].

There appears to be heterogeneity in tissue residency programs across different cancers types. For example, we identified *ZNF683* (HOBIT), a central mediator of lymphocyte tissue-residency in mice [65], as an Exh marker with relatively higher abundances in CD8 and NK Exh compared to their respective Res cells. This is in contrast to NSCLC where relatively high expression has been shown in resident T cells [7, 65]. These were confirmed in studies where *ZNF683* expression was higher in Res cells in NSCLC compared to hepatocellular carcinoma and CRC [6–8]. While we identified the NR4A family genes as Res signature genes, the NR4A family have been previously shown to be linked to CD8 T cell exhaustion [66] and shown to limit the function of chimeric antigen receptor (CAR) T cells in solid tumours [67]. Marquardt et al reported the up-regulation of *ZNF331* within the CD69^+^CD49a^+^CD103^+^ resident NK cells from normal lung tissue compared to CD69^-^ cells [68], and we identified this gene as one of our CD8_Res genes. While they also observed higher expression of *IRF4* in the resident NK cells, we removed this gene from our list due to a particularly high correlation with both Res and Exh programs in NK cells. Notably, we here identified *IRF8* as a TF involved in Res programs.

The unsupervised trajectory analysis suggested that exhausted CD8 T cells drive from both resident cells and circulating CD8 T cells. This is in line with the reported diversity in the origins of exhausted CD8 T cells [5]. Zhang et al observed a developmental link between effector memory T (T_em_) cells and exhausted T cells in CRC patients [8], while in NSCLC, Guo et al showed a direct link between T_rm_ and exhausted cells [7]. Li et al showed that the dysfunctional (or exhausted) cells in melanoma patients drive from transitional (early effector GZMK^+^) T cells through a local differentiation process, without any evidence of exhaustion in peripheral blood [19], while Huang et al analysed exhausted CD8 T cells in peripheral blood of patients with melanoma that were treated by anti-PD1 therapy, and had T cell clones in common with the tumour [20]. Interestingly, the obtained trajectory of the NK and CD8 Res cells in our results passed through an intermediate sub-population with co-expression of *GZMK,* naïve cell markers and some inhibitory receptors. Similarly, Galletti et al recently reported two subsets of CCR7+ CD8 T cells, one of which expresses *GZMK* along with *PDCD1* and *TIGIT,* which they termed T precursor exhausted-like cells, which commit to a terminally dysfunctional state [69]. Therefore, this intermediate sub-population of NK cells may represent the NK precursor exhausted cells.

Based on our mutation analysis, the NK_Exh_Bulk signature was associated with the highest number of gene mutations. This suggests that tumour mutation burden and related neoantigen expression may drive tumour inflammation and consequently impact NK cell activation and exhaustion. Another interpretation is that in the presence of highly activated NK cells, tumour cells may undergo positive selection for cells with mutations resulting in decreased immunogenicity to these cytotoxic lymphocytes as an immune surveillance escape strategy [70]. This is also supported by our findings of gene mutations that were predictive of NK_Exh_Bulk (e.g. genes associated with antigen presentations). Another predictive gene of NK_Exh_Bulk was *POLD1,* mutations of which have recently received attention as biomarkers for cancer immunotherapy due its crucial role in DNA replication and repair [71, 72]. We also found several less appreciated genes in this context, such as *TGM6,* a transglutaminase, which similar to other members of TGMs, such as *TGM2,* performs post-translational modification in a Ca^2+^ dependent manner, and may play a role in cancer [73]. We also identified calcium ion transmembrane transport as one of the significant GO terms associated with NK_Exh_Bulk genes, and interestingly, not only intracellular Ca^2+^ concentration is required for cancer cell proliferation and apoptosis, but also, the efficacy of the cytotoxic T cell and NK cell function is dependent on Ca^2+^ [74].

GO enrichment of the mutations associated with NK_Exh_Bulk also suggested the importance of several actin filament-based and cytoskeletal processes, which are much less appreciated in the context of tumour immune infiltration. Solid tumours have been shown to undergo massive remodelling after attack by cytotoxic T and NK cells; this remodelling not only impacts tumour cell recognition by lymphocytes, but also influences immune cell activation [75]. We also noted that some of our Exh signature genes (such as *MYO7A* and *AFAP1L2,* shared by CD8 and NK cells) were those associated with actin and myosin-based processes. Remodelling of actin filaments and myosin contractibility are required for the formation of immunological synapse as well as transportation and secretion of cytotoxic granule in T and NK cells [76], and increased expression of some actin modulating genes was also reported in rapidly cycling NK cells which exhibited reduced cytotoxicity [62].

Mutation and survival analyses suggested important associations of Exh/Res programs with oxidative phosphorylation (or hypoxia), DDR, TP53 and cell cycle processes. Oxidative stress can cause DNA damage, and if DNA damage response (DDR) is not effective, it leads to tumour initiation and promotion in a tumour favourable microenvironment [77]. We observed negative correlations between Exh/Res scores and DDR, suggesting that a defective DDR programs could lead to more neo-antigens and therefore more immune infiltration. However, our also analysis indicated that samples with lower CD8NK_Exh_Bulk and low DDR have relatively poor survival outcome compared to those with high CD8NK_Exh_Bulk and high DDR scores, which may suggest that some activity of DDR could be beneficial when combined with exhaustion programs from CD8 and NK cells. This is consistent with reports that suggest defective DDR not only can stimulate immune response, but also can lead to immunosuppression [78]. Further, our results also showed that active TP53 signalling, an important mediator of the DDR, when combined with high CD8NK_Res_Bulk programs is suggestive of a relatively poorer prognosis. This is in line with previous studies, reporting that mutations in TP53 is associated with higher immunogenic activity in a number of cancer types [79], which may be associated with a relatively higher Exh programs as a result. Altogether, these results suggest that depending on the relative Exh and Res programs in tumour microenvironment, defects in DDR and TP53 pathways could lead to a better or worse survival outcome.

In both single cell and bulk tumour data, we observed positive correlations between TGF-β-related signature scores and Res/Exh scores, and this correlation was higher for Res scores compared to Exh scores. These results are not surprising considering the importance of TGF-β in the development of resident T cells [29, 32]. In this study, we took this a step further and demonstrated how combining TGF-β signalling scores with NK_Res_Bulk or NK_Exh_Bulk scores could predict survival outcome. We also, for the first time, showed the importance of the combined use of NK_Res_Bulk and NK_Exh_Bulk signatures for predicting survival outcome, independent of known CRC clinical factors (e.g. age, stage, MSI and CMS). This is particularly important considering that MSI status alone is not sufficient in predicting response to immunotherapy or survival outcome [2, 48], ~13% of CRC patients cannot be assigned to any of the CMS subtypes and almost all CMS subtypes include a mixture of samples with both poor and good prognosis, the exception being CMS4 and relapsed CMS1, which are associated with poorer survival outcomes [80]. By taking into account that: (i) samples with lower TGF-β signalling and lower NK_Res_Bulk are associated with better survival, while samples with lower NK_Exh_Buk and higher TGF-β signalling show poorer outcome, (ii) TGF-β has been associated with T cell exclusion in several cancer types and targeting TGF-β has been shown to be beneficial for response to ICIs [36], and (iii) we observed better survival outcome for patients with high NK_Exh_Bulk and low NK_Res_Bulk programs (compared to those with low NK_Exh_Bulk and NK_high Res_Bulk), it is tempting to speculate that part of the suppressive biology associated with TGF-β signalling is by promoting NK residency.

In this study, we suggested a better survival outcome for patients with higher Exh to Res programs (specifically for NK and combined NK and CD8 signatures) by defining these programs via non-overlapping gene signatures. However, several previous studies reported associations between CD8 Res cells and better response to ICBs and improved survival outcome [24–26]. While some of these could be due to in cancer types and heterogeneity in Res cell populations, it is also at least partially due to the use of a non-specific signature or a mixed Exh and Res gene sets in these studies, and as noted above, there is a strong degree of overlap between T cell activation and Exhaustion markers which are difficult to distinguish in the absence of time course data. For example, many studies defined Res cells using *ITGAE* (CD103) [25, 26]. We removed this gene based on the results of the canonical correlation analysis on the initial Res and Exh signatures in NK and CD8, which means that the presence of this gene was associated with both Res and Exh programs. Duhen et al identified a subset of CD8 TILs with T_rm_ phenotype with co-expression of *ITGAE* and *ENTPD1* (CD39), which were also enriched for exhausted markers *(PDCD1, CTLA4,* and *HAVCR2)* and had high expression of *CD69;* they showed that these cells were associated with high tumour-reactivity and better prognosis in head and neck cancer patients [81]. Therefore, high expression of *ITGAE* and better survival outcome may not be specifically attributed to Res cells. Similarly, we removed *ITGA1* (CD49a) from the CD8 Res signature for the same reason; however, *ITGA1* was a good Res marker for CD4 cells. Another example is the T_rm_ signature reported by Byrne et al [24], which includes a mix of Res (e.g. *CD69)* and even more Exh markers *(HAVCR2, PDCD1, TIGIT, LAYN,* etc). The current study not only introduces novel distinct Exh and Res programs in CRC T cells and in particular in NK cells, but also shows the importance of such signatures and their combinations with known genomics and molecular information in stratification of samples to predict survival outcome and potentially their response to immunotherapy.

## Methods

All analyses for this study were performed using R version 4.0.2, with Bioconductor version 3.11 [82]. A list of all the packages and tools used in this study with their versions and citations are given in Supp. Table S7.

### Deriving initial exhaustion and residency signatures

We used high-quality single cell data (Smart-seq2) from Zhang et al (2020) [21] to obtain gene expression signatures. Transcript per million (TPM)-like data were obtained from GSE146771 on 9^th^ of Jun 2020, with data extracted for CD8, CD4 and NK cells using pre-defined annotations from the meta-data, and analysed each cell type separately.

In each case, we started with a selection of canonical markers associated with residency (Res) and exhaustion (Exh) in each of the cell types. The Res canonical markers consisted of *CD69, ITGAE* (CD103), *ITGA1,* and *RGS1* for CD8_Res and CD4_Res, and we additionally included *ZNF683* (Hobit) for NK_Res. The Exh markers included: *HAVCR2* (Tim-3), *PDCD1* (PD-1), *CTLA4, LAYN, CXCL13* for CD8_Exh; the same genes plus *LAG3* for CD4_Exh, and; *HAVCR2, PDCD1, CTLA4, TIGIT,* and *LAG3* for NK_Exh.

Next, we curated an extensive list of genes reported in the literature to be associated with residency or exhaustion across different species (human & mouse), as well as conditions (infectious disease, cancer and normal tissue) (Supp. Table S8). For each cell type, we then calculated the Spearman’s correlation (ρ) between candidate Exh and Res markers and the list of canonical markers for that cell type. For NK cells, genes with ρ > 0.35 with any of the canonical Res signatures were considered to be Res genes, and those with ρ > 0.25 with canonical Exh genes were counted as Exh markers. These thresholds were slightly different for CD4 cells due to differences in their gene expression distributions (ρ > 0.3 for CD4 Res and CD4 Exh). After removing genes that were overlapping between the Res and Exh gene list, we performed a sparse canonical correlation analysis (CCA; using PMA R package) and removed genes with a coefficient < −0.1 in the first component in order to reduce the correlation between the Res and Exh sets. We refer to the resultant gene sets as the ‘initial’ Res and Exh genes; these were different for each cell subset, although there some genes were overlapping across subsets.

### Deriving the final exhaustion and residency signatures

Using the initial Res and Exh signatures and the singscore R package, we scored the Zhang Smart-seq2 NK, CD4 and CD8 cells. We then stratified cells based on their Res and Exh scores and performed differential expression (DE) analyses between cells with high Res/low Exh scores and those with high Exh and low Res scores. We used the Seurat R package to perform DEanalyses with two different approaches: (i) Wilcoxon Rank Sum test, and; (ii) the hurdle model from the MAST package. We retained genes that were considered as differentially expressed using either approach (adjusted p-value < 0.05 and logFC > 0.3), and further refined this list by comparing transcript abundance percentiles for Res vs Exh vs circulating blood cells. Specifically, we retained Exh genes whose 65^th^ %ile abundance in Exh cells was higher than the 90^th^ %ile expression in both Res and blood cells. Similar comparisons were performed to refine Res genes, and a 75^th^ %ile threshold was used when defining the NK_Exh signature.

### Analysis of single cell data sets

For each of the single cell data sets used in this study, we considered each cell subset independently, and therefore, the data filtration, clustering and dimensionality reductions were also performed using slightly different parameters in each subset of a given dataset. We refer readers to the code available on Github for details on parameter settings. For all raw single cell data, we performed the SCTransform method from the Seurat package using the top 5000 variable genes, regressing on the percent of mitochondrial genes. We also used the Seurat package to perform clustering, dimensionality reduction, and DE analysis. The slingshot R package was used to perform the trajectory analysis. For all single cell data sets, we used the singscore package to score single cells. Blend plots were generated using the Seurat visualisation functions with single-cell scores obtained from singscore.

Processed Zhang et al single cell RNA-seq and metadata for both Smart-seq2 and 10X data were obtained from GSE146771 on the 9^th^ of June 2020 [21]. No further normalisation or processing was performed. The de Vries et al single cell RNA-seq data as well as cell annotation information were received from the authors 23^rd^ of July 2020 [41]; we included two cell groups annotated as proliferative cells and unclassified cells when analysing the data for each of the NK, CD4, and CD8 T cells separately. The raw single cell RNA-seq data from the human gut atlas by James et al were downloaded on 9^th^ of June 2020 from the Gut Cell Atlas (https://www.gutcellatlas.org/) [42]. We performed batch correction to remove donor differences using the integration approach provided in the Seurat package.

Raw single cell RNA-seq data from de Andrade et al were downloaded for one melanoma patients (CY129; with a relatively large number of infiltrating NK cells) on 15^th^ of Jun 2020 from GSE139249 [43]. We analysed these data in two steps: first, we ran the Seurat pipeline and normalised the data in order to identify potential contaminating cells based on the expression of marker genes for cells other than NK cells. To be consistent with the original paper, we used the same marker genes for filtration *(CD3D, CD3E,* and *CD3G* for T cells; *IGHG1, IGHG2,* and *JCHAIN* for B cells; *LYZ* for macrophages; and *MLANA* for melanoma cells). Then we filtered these from the original raw data. From 11368 cells, we retained 4267 with no expression for the above markers. We further filtered cells to remove those with: > 20% mitochondrial genes, number of RNA counts ≥ 25,000, or; a number of features ≤ 500, which resulted in 4195 cells. Next, we used the Seurat package to analyse these data using similar settings provided above; we performed SCTransform for normalisation, used the first 20 PCs for identifying the k-nearest neighbours of each cell and performed dimensional reduction. For running UMAP, we also specified n.neighbors = 50, min.dist = 0.4.

To obtain proliferative signatures in the Exh single cells from Zhang et al Smart-seq data, we stratified cells based on their exhaustion and proliferation scores, and then used Seurat package and performed DE analysis between cells with high Exh and low proliferation scores and those with high Exh and high proliferations scores.

### Refinement of the signatures based on bulk cell line and laser capture micro-dissection data

The RNA-seq count data and meta-data for all cell lines from the cancer cell line encyclopedia (CCLE) were downloaded from the Broad Institute Data Portal (https://portals.broadinstitute.org/ccle/) [83] on the 7^th^ of Feb 2020. We downloaded gene annotation information (e.g. gene length) from Gencode (v19) and used the edgeR Bioconductor package to generate a DGEList object in R. We then filtered the pan-cancer data to only retain genes with cpm > 1 in at least 5 cell lines, performed TMM normalisation and calculated the RPKM values. The data were then subset to only visualise the expression of the marker genes in the CRC cell lines (Figure 2). Comparing against these data, we considered Exh and Res markers to pass the “bulk tumour threshold” if they had median logRPKM expression ≤ 0.

We also downloaded processed microarray data (log2 RMA normalised) from high purity CRC samples collected with laser capture microdissection (LCM) by Tsukamoto et al [45] (GSE21510) using the GEOquery R package in September 2020. Probes which mapped to more than one Entrez ID were excluded, and for probes mapping to the same gene, we considered the average expression value. In these data, we considered Exh and Res markers to pass the “bulk tumour threshold” if they had median logRPKM expression ≤ median expression of all genes in the data. Finally, if a given gene passes the “bulk tumour threshold” in CCLE or LCM, we considered that marker to be suitable for use in bulk tumour samples. The scores from these markers have “Bulk” prefix in their names. We note that that abundance data for *MARCH3* did not exist within the CCLE data, and 14 genes *(KRT86, APOBEC3H, TTC24, MIR4435-2HG, HNRNPA1L2, HLA-DRB5, HLA-DRB1, KLRC2, ADGRE5, HSPA1A, HSPA1B, XCL2, HSPA8, NOTO)* were not present within the LCM microarray data, and therefore, could not be validated against these data sets.

### Analysis of the transcriptomics data from bulk tumour samples

Raw counts for TCGA COAD harmonized RNA-seq data [84] were downloaded using the TCGAbiolinks R package. We only considered protein coding genes and filtered to remove lowly-expressed genes (genes with count < 15 across 90% of the samples, applied in cancer and normal samples separately) and data were normalized using RUV-III.

Processed data from Marisa et al were downloaded from GSE39582 on 18^th^ of March 2019 using the GEOquery R package, and a SummarizedExperiment object was generated using the SummarizedExperiment Bioconductor package. Consensus molecular subtyping in both data sets was performed using the CMScaller R package.

### Single-cell and single-sample scoring, and molecular signatures

We used the singscore Bioconductor package to score both single-cells and single samples where relevant in this study. We only used signatures with known or expected up-regulated genes, and did not centre scores, such that scores range between 0-1, where 0.25 corresponds to a median signature gene rank in the 25^th^ %ile by abundance, and 0.75 corresponds to a median signature gene rank in the 75^th^ %ile by abundance. A list of tumour related signatures that were used to score tumour data and perform survival analysis, as well as signatures used to score single cell data are given in Supp. Table S9.

### TFs and GPCRs

A transcription factor (TF) list was obtained from the Human Protein Atlas (HPA), by searching for “Transcription factor” in *gene description* or *protein class* fields, identifying 1549 TFs. We generated a list of genes that were GPCRs or those associated with GPCR signalling pathway from several sources, including the HPA (searching for “coupled” in *gene description* and *protein class* fields), GO terms (obtained from QuickGO), and uniprot (searching for “gpcr” under the *Type* field), resulting in 2456 genes (Supp. Table S10).

### Identification of genomic mutations associated with residency and exhaustion

The TCGA mutation data (maf format) were downloaded using the TCGAbiolinks package on 15^th^ of Oct 2018. We filtered the mutation data for those with low or modifier impacts, and retained mutations with moderate or high impacts. Several mutations in the same gene for the same sample were annotated as multi-hit (MHT), and duplicated records were removed to retain one record of mutation in a given gene within individual samples. Using this data, we defined the mutation load as the total number of mutated genes in a given sample.

We then subset the mutation data to keep genes mutated within at least 10 patient samples. For each of the Exh and Res scores (from the Bulk signatures) for different cell types, we used the tidyverse and broom R packages to generate several linear models with score as output and presence or absence of individual gene mutations as input. The obtained p-values were then adjusted using Benjamini-Hochberg (BH) method for multiple hypothesis testing. Mutations with adjusted p-values < 0.01 were considered to be predictive of scores.

To perform GO analysis on the predictive genes of NK_Exh_Bulk, we first subset the significant predictive genes to those with adjusted p-value < 0.0001 (*N_Genes_* = 1027), and then used the goana function from limma package. From GO biological processes (BP) with FDR < 0.01, we removed terms with <= 20 or >= 1500 genes. Finally, we used the rrvgo R package to calculate semantic similarities between GO terms and visualise the GO parent terms.

For the NK_Exh and NK_Res (Bulk) signatures, we also performed elastic net regression using the glmnet R package. We performed 10-fold cross validation using the cv.glmnet function for 100 lambda values between 0.01 and 10^10^ to obtain the optimal lambda value, and used that in the final elastic net model. The optimal lambda value for NK_Exh_Bulk was 0.0175 and for NK_Res_Bulk was 0.01. Inputs to the elastic net regression included all genes with showed significant associations against NK scores in the linear model (the above analysis), as well as additional parameters such as: mutational load; stage; MSI status, and; CMS cluster. To perform elastic net regression, we removed samples with NA values for the four additional parameters, and again filtered for genes that had mutations in at least 10 samples. This yielded a final data set with 3238 genes and 305 samples (which were used to generate boxplots in the Supp. Figures S10–S12). We then considered parameters with beta values > 0 or < 0 to be predictive of NK scores.

### Survival analysis

In each of the TCGA and Marisa data sets, for a given survival analysis (i.e. OS, PFI, and RFS), we first performed lasso regression using the glmnet R package to select Cox model covariates. Variables tested were age, stage, MSI status, and CMS subtype. We then selected variables with non-zero coefficients using minimum lambda values obtained from crossvalidation (using the cv.glmnet function with the parameter family = “cox”), and used those as covariates when performing Cox multivariate analysis with scores or score pairs. These covariates were different in the two data sets and from one survival type to the other: for TCGA OS and PFI: age, stage, MSI, and CMS; for Marisa RFS: age, stage, MMR, and CMS, and; for Marisa OS: age, stage, and CMS.

To examine the associations between different scores and survival outcome (by including relevant covariates), we first examined each individual Exh and Res scores by dividing samples into two groups of high and low scores based on median values. We also considered median scores when stratifying samples based on score pairs (including Exh and Res signature scores as well as tumour related signature scores). Finally, in an attempt to compare samples with more distinct Exh and Res programs for each of the immune cell types in cancer, we subset the data to only retain samples with high Exh & low Res scores and those with low Exh & high Res scores, and then generated Cox models comparing the survival outcomes of these two groups while accounting for relevant covariates.

## Code availability

The R Markdown reports, R scripts, and small data dependencies reproducing the results of this study are on Gitlab (https://gitlab.com/huntington-immuno-lab/foroutan_resexh_crc).

## Acknowledgements

This work was supported by project grants from the National Health and Medical Research Council (NHMRC) of Australia (1124907, 1124784, 1049407, 1066770, 1057852, 1027472) and a fellowship (1124788) to N.D.H. N.D.H. is a recipient of Research Grants from the Harry J Lloyd Charitable Trust (USA), Melanoma Research Alliance (USA), Tour de Cure (AUS), the Ian Potter Foundation (AUS), Cancer Council of Victoria (1145730) and a CLIP grant from Cancer Research Institute (USA).

The results shown in this study are in part based upon data generated by the TCGA Research Network: https://www.cancer.gov/tcga, and the CCLE data obtained from the Broad Institute: https://portals.broadinstitute.org/ccle. The authors thank Dr Miranda and Dr de Vries from Leiden University Medical Center for providing single cell data (de Vries et al (2020), Gut) used in this study.

## Authorship Contributions

MF and NDH conceived and designed the study; MF, RM, JC, and TS performed the data analysis; MF, JC and NDH interpreted all data. All authors wrote and read the paper.

## Supplementary Figures of the paper entitled

**Figure S1.**
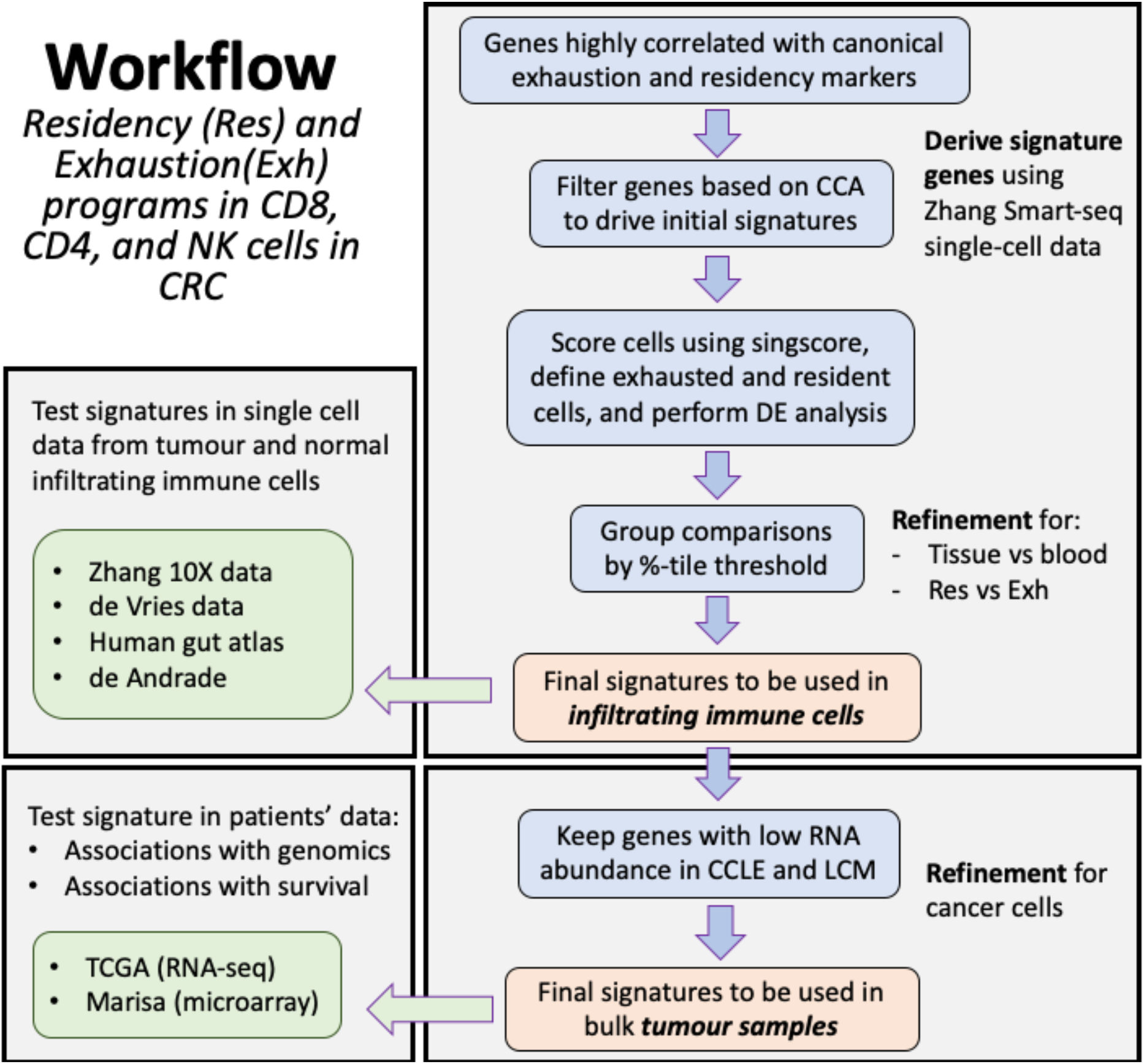
Workflow of the analyses performed in this study.

**Figure S2.**
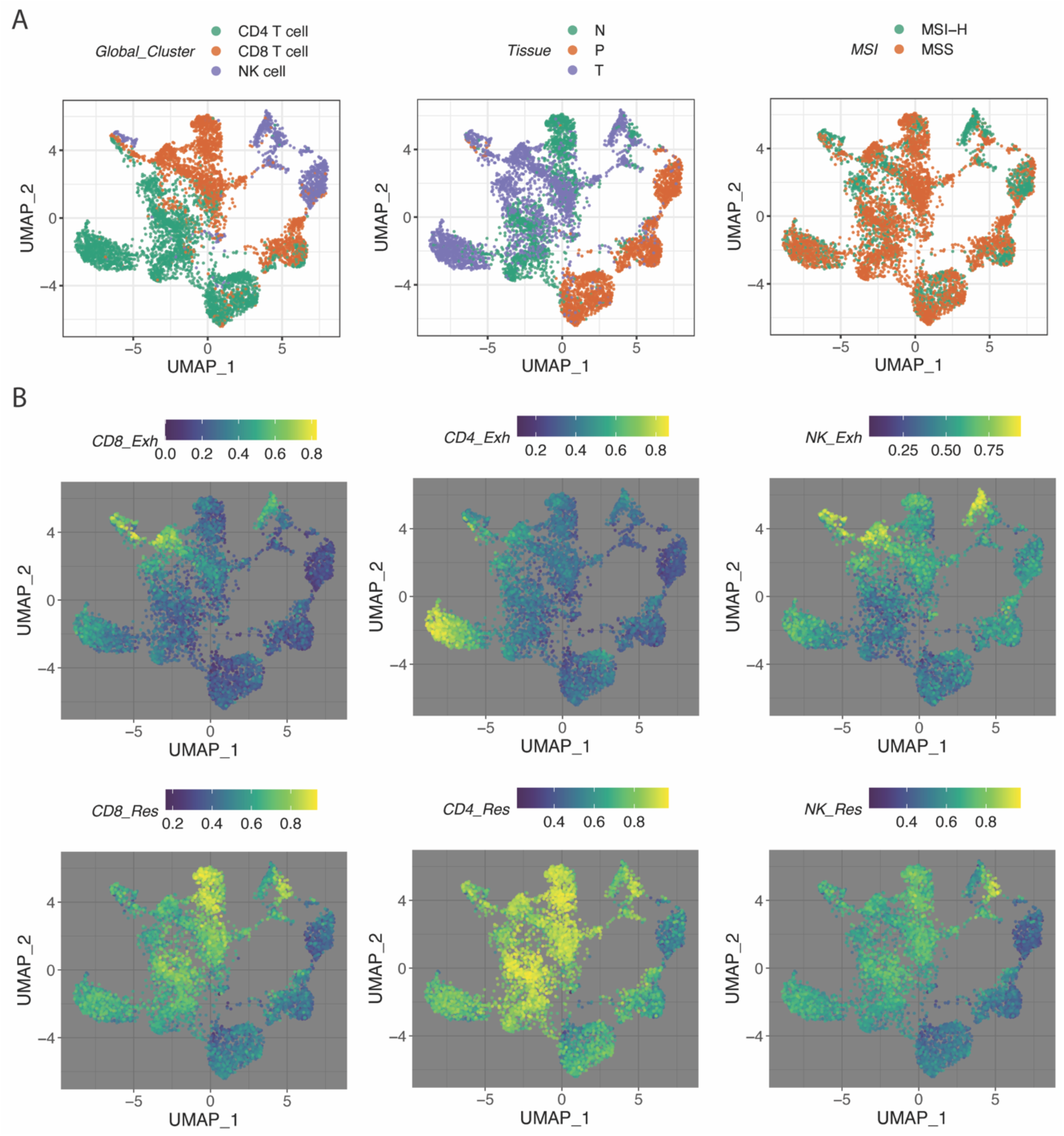
UMAP plots of the CD8, CD4 and NK cells from Zhang et al Smart-seq2 single cell data, coloured by cell annotations (top row), or Exh and Res signature scores for each of the cell types. N = normal, T = Tumour, P = Peripheral blood.

**Figure S3.**
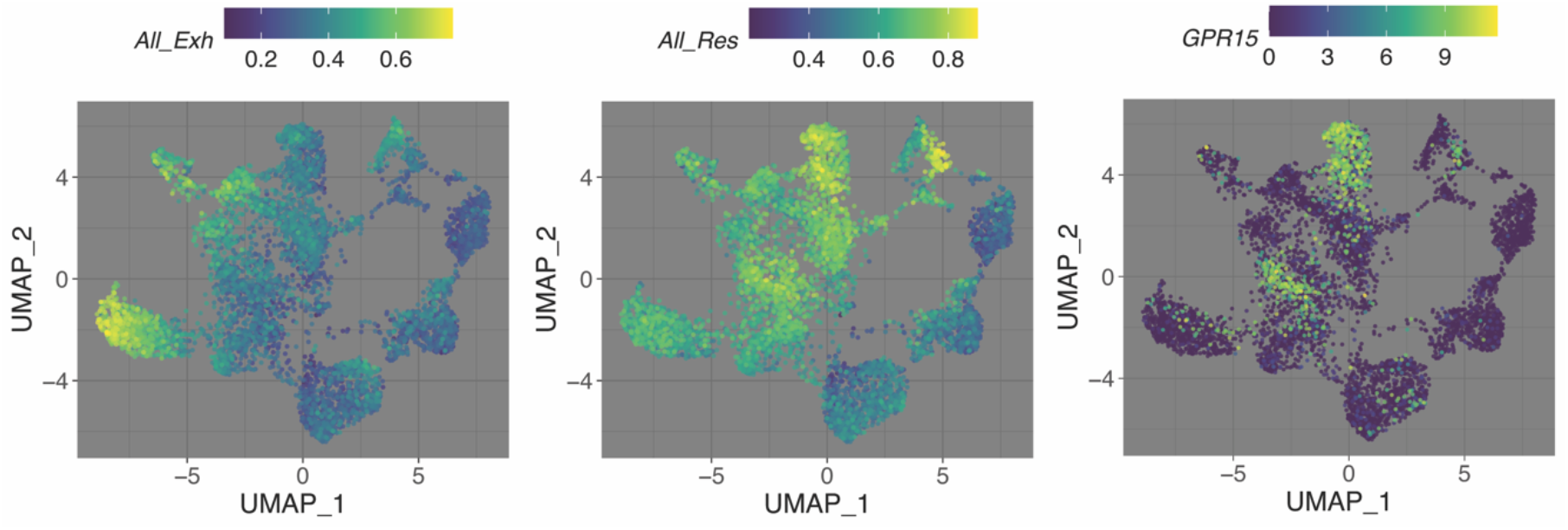
UMAP plot of the of the CD8, CD4 and NK cells from Zhang et al Smart-seq2 single cell data, coloured by Exh and Res scores from all three signature sets (left and middle figures), or by the expression of GPR15, one of the Res markers.

**Figure S4.**
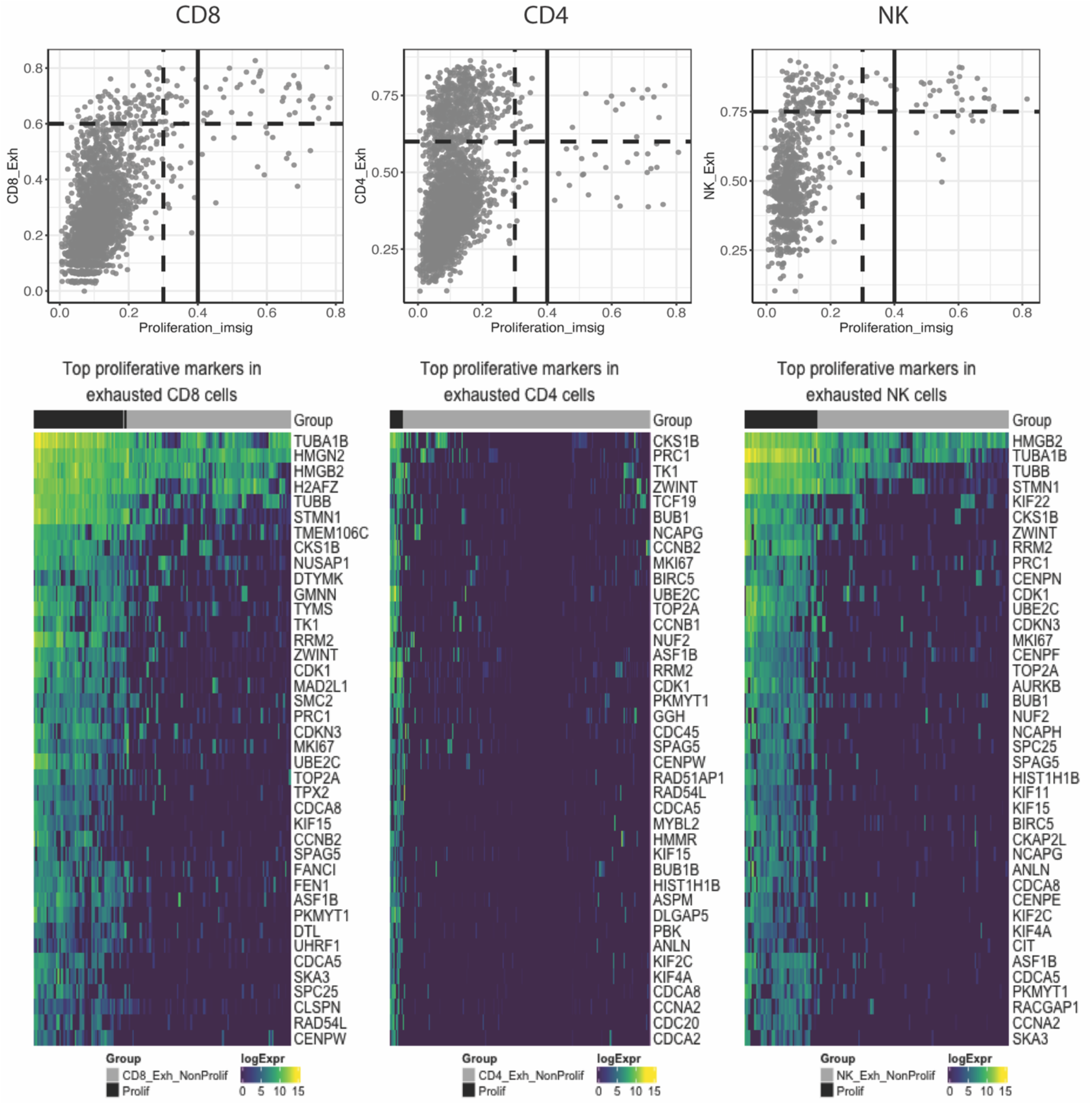
Exhausted immune cells showing the two sub-populations of proliferative and nonproliferative cells. In order to understand the heterogeneity across the exhausted population of T and NK cells, we stratified samples based on their exhaustion and proliferation scores, and obtained markers that were highly expressed in proliferative exhausted cells compared to non-proliferative exhausted cells (Suppl Table S3). Among these, 138 genes were common across the three cell types, of which only ECT2 was involved in GPCR signalling pathway and TCF19 transcription factor (TF) were shared across the three cell types. CALM2 was another GPCR that was shared by proliferative CD4 and NK exhausted cells. TFs that were common in exactly two cell types included: FOXM1, TFDP1, and DNMT1 in CD8 and CD4; E2F2 in CD8 and NK; and MXD3 in CD4 and NK cells.

**Figure S5.**
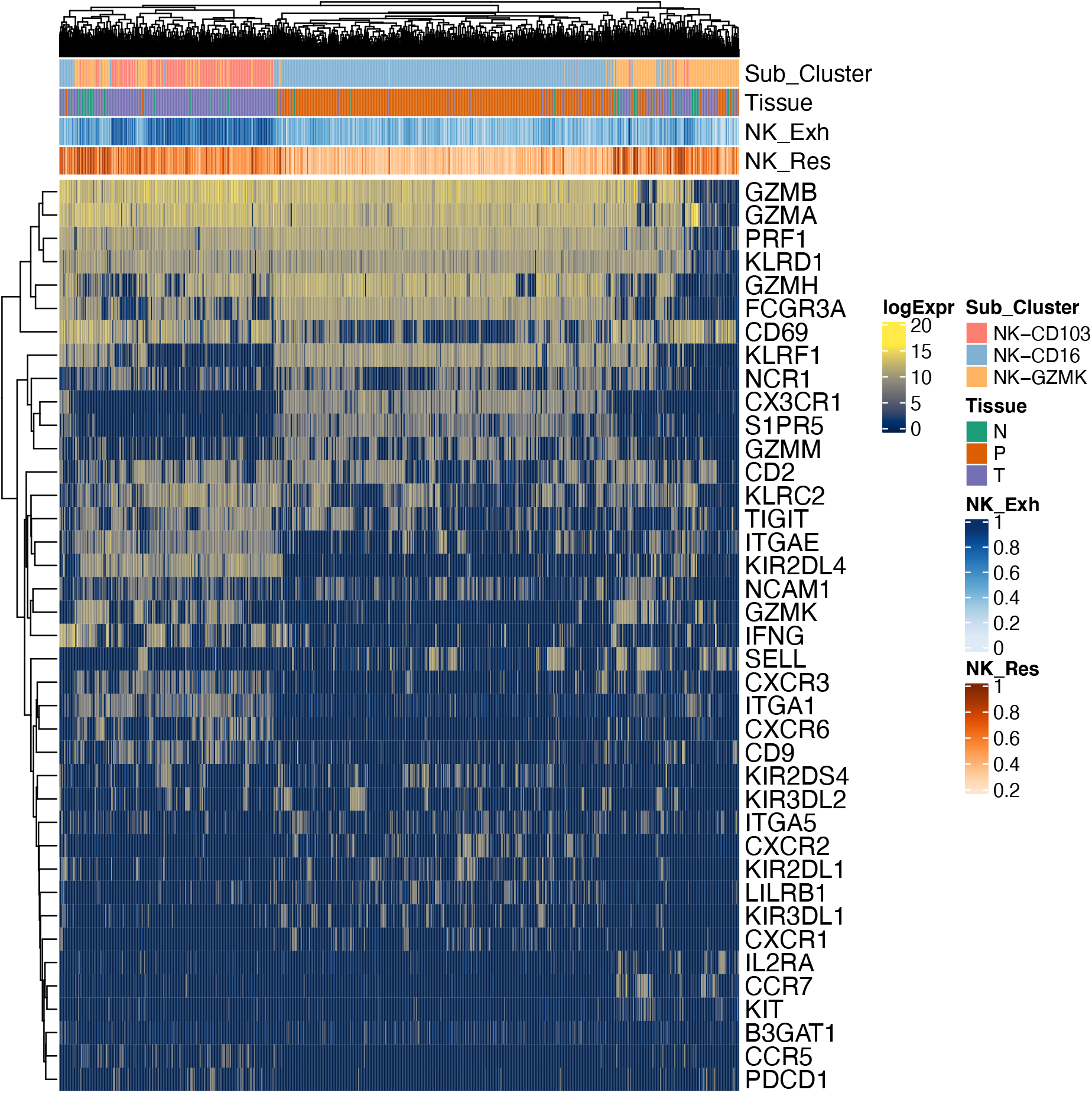
Expression of selected genes in the NK cells from Zhang et al smart-seq2 data. Heatmap of gene expression for selected genes (from Frued et al) in all the NK cells from Zhang et al Smart-seq2 data, including those from peripheral blood (P), tumour (T), and adjacent normal tissue (N).

**Figure S6.**
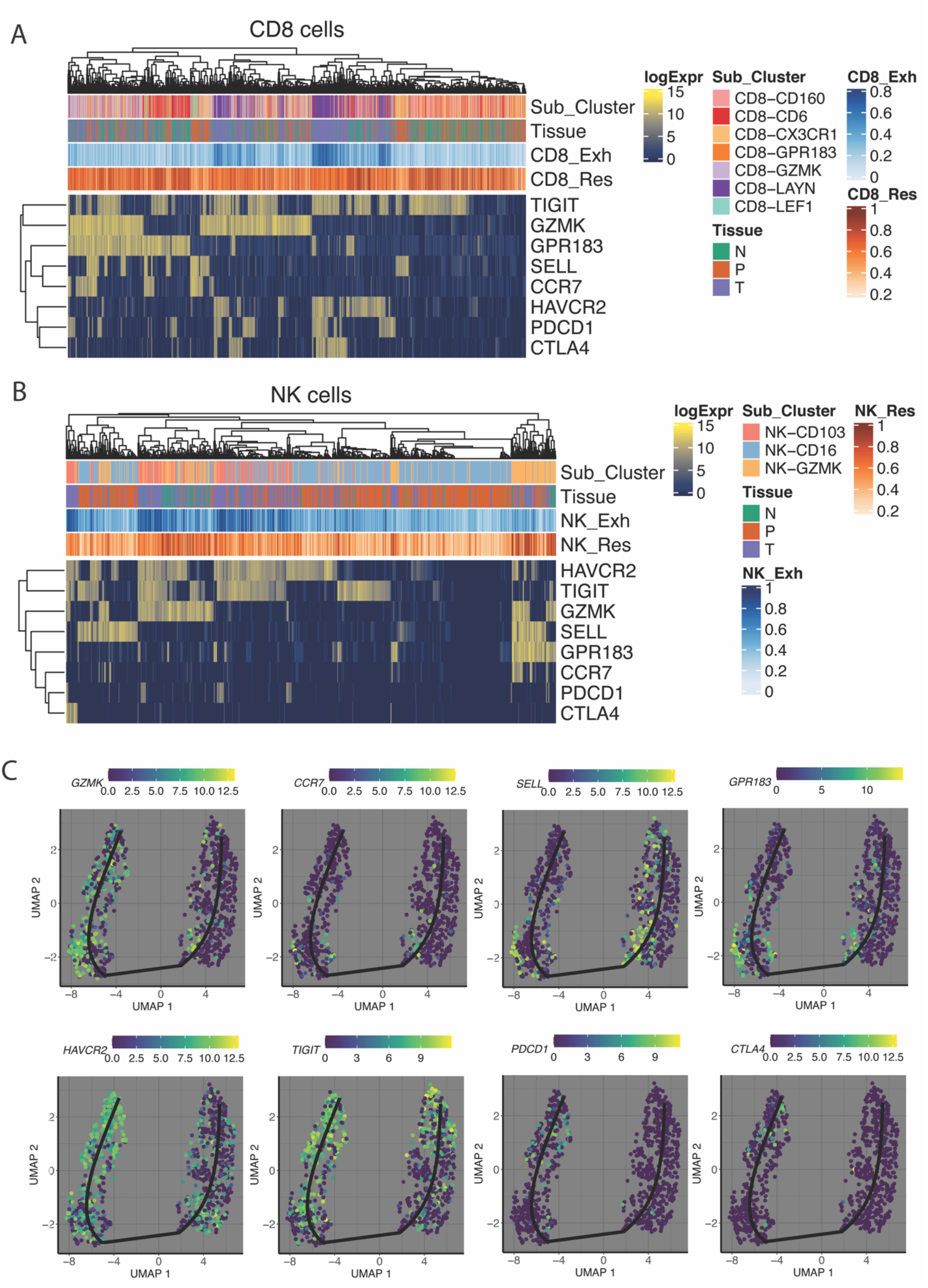
Expression of selected genes in the CD8 (A), and NK cells (B) in Zhang et al Smart-seq data. We observed co-expression of naïve genes (SELL, CCR7, and GPR183) with inhibitory receptors (at least one of the CTLA-4, PDCD1, TIGIT, and HAVCR2) in a sub-population of GZMK+ cells. (C) Trajectory analysis of the NK cells, coloured by the expression of the same genes, which highlights a sub-population of cells co-expressing some of these genes in the intermediate stage of the trajectory.

**Figure S7.**
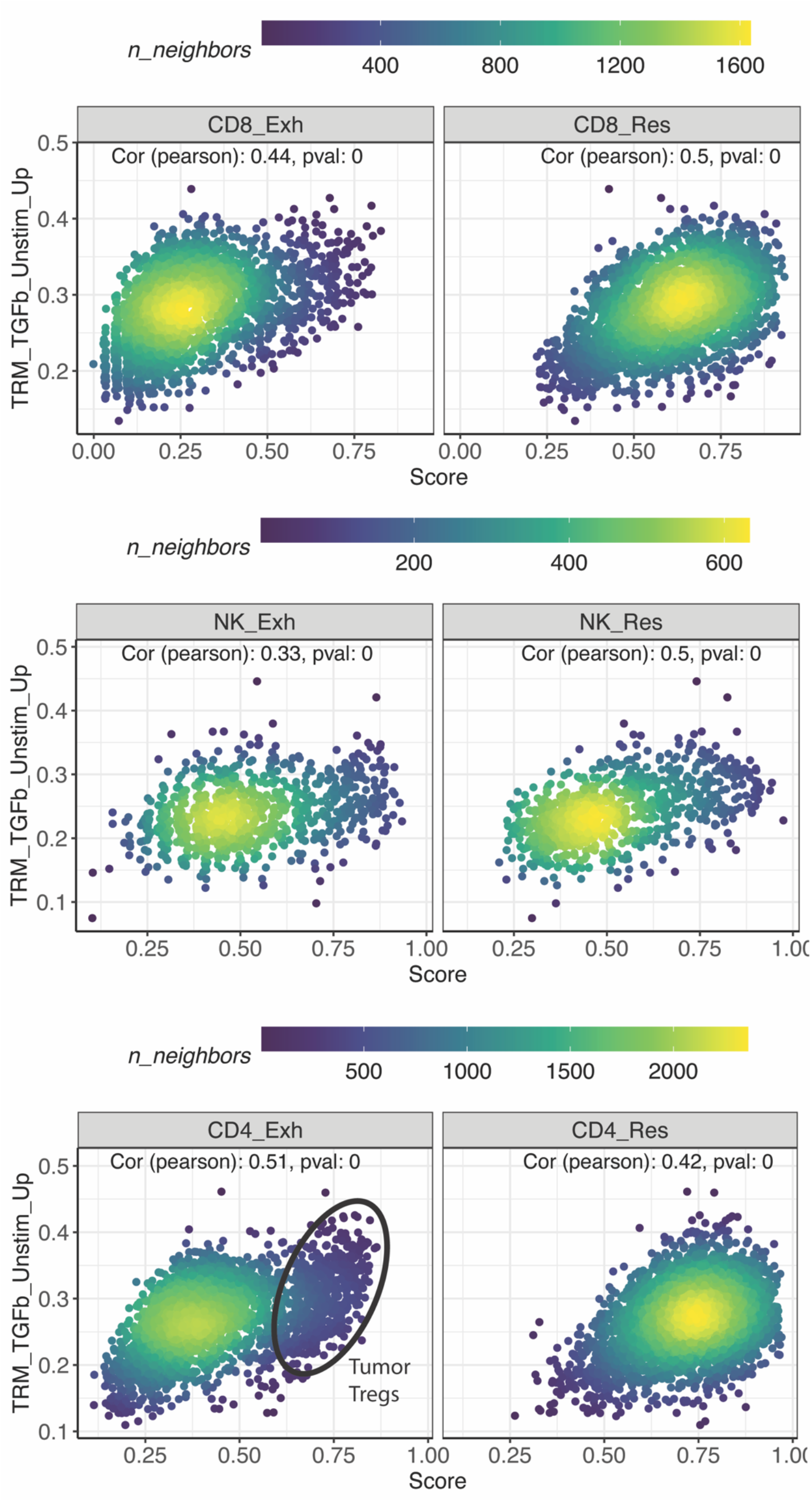
Association between Exh and Res signature scores and TGF-β signatures from tissue resident memory cells (from Nath et al) in each of the CD8, CD4 and NK cells from Zhang et al Smart-seq2 data.

**Figure S8.**
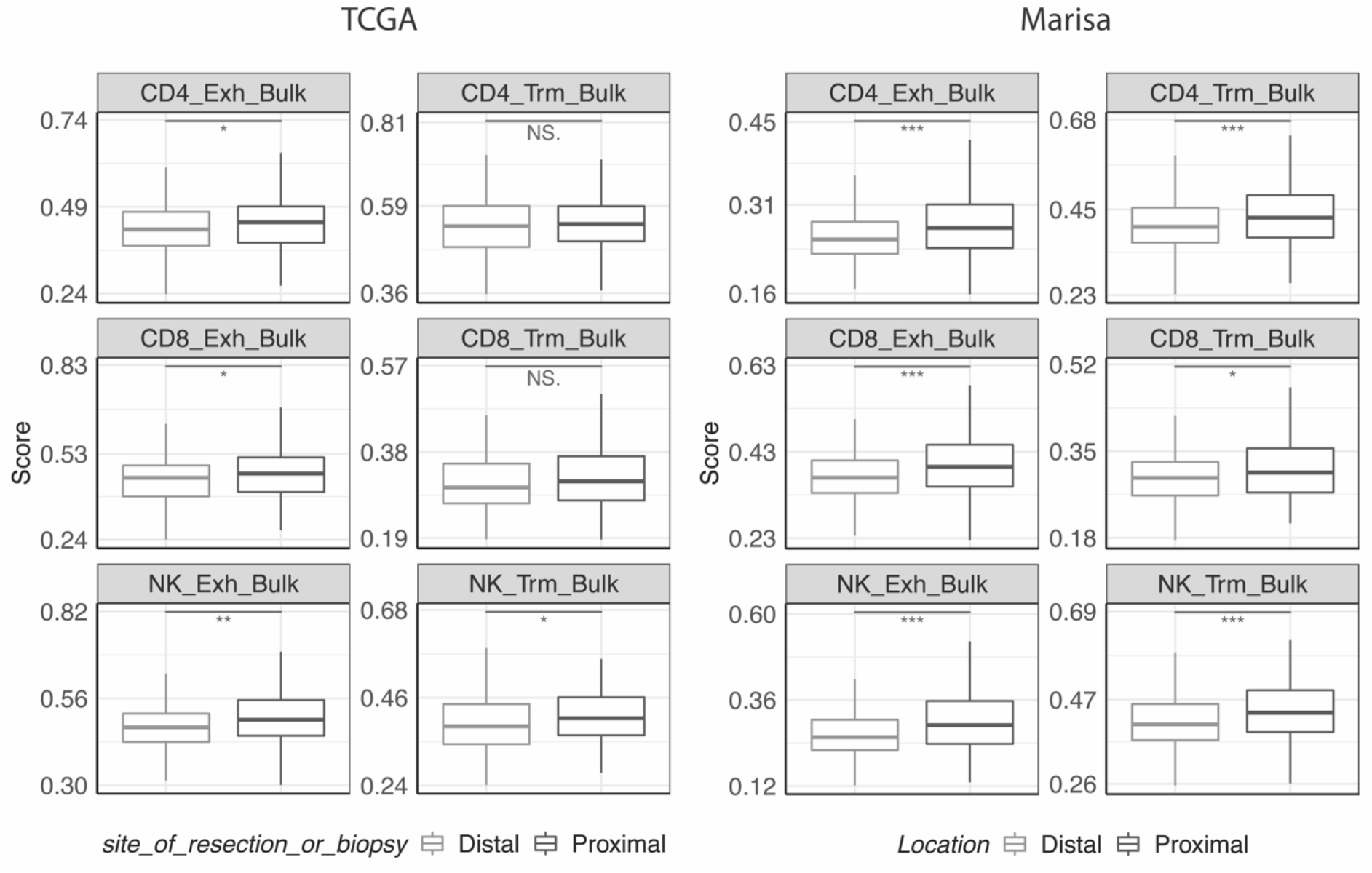
Exhaustion and Res scores across tumour samples from different locations (proximal and distal) in the TCGA and Marisa data.

**Figure S9.**
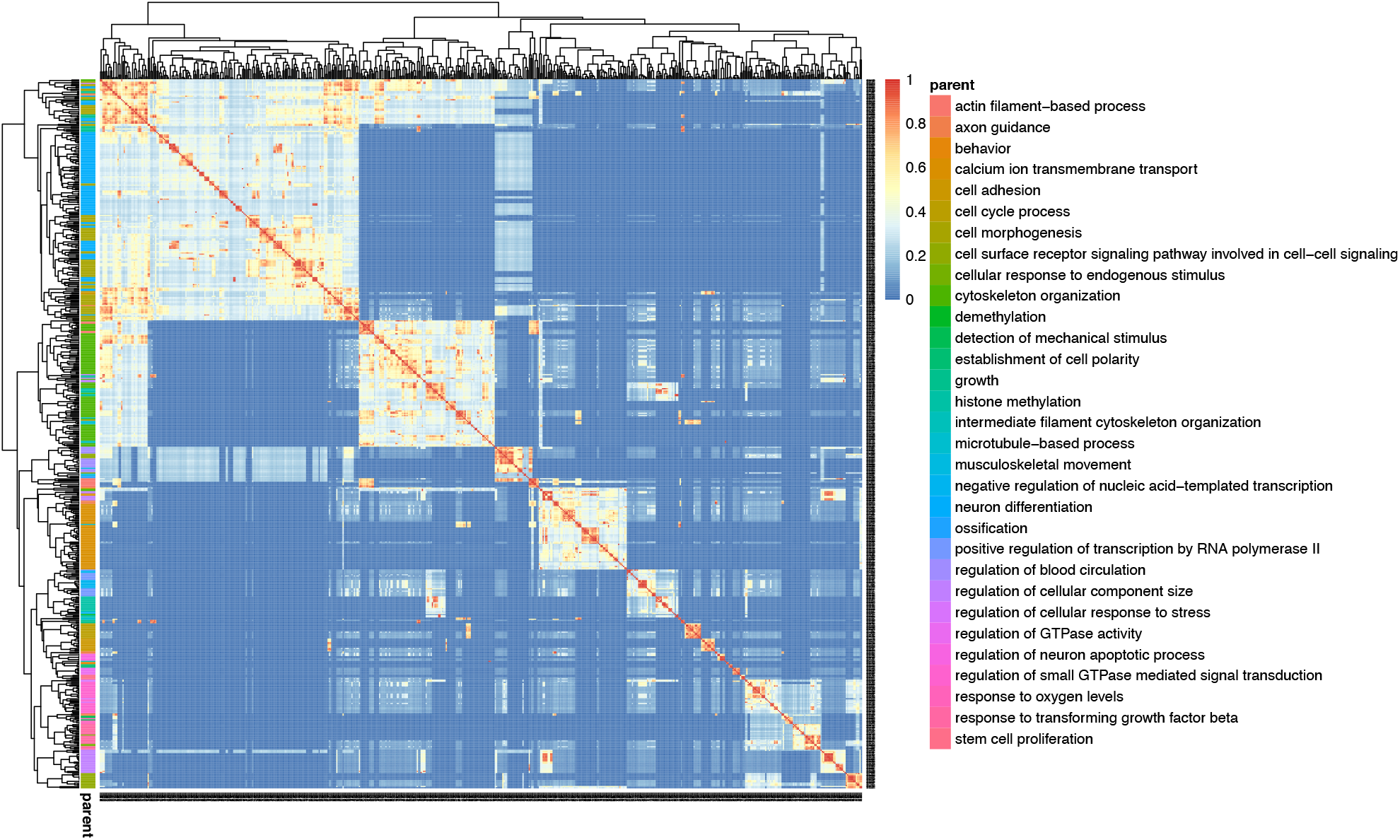
Similarity matrix of top significant GO terms using top selected genes whose mutations were significantly associated with NK_Exh_Bulk. The heatmap summarises 606 terms that are annotated with their parent terms. Please see Supp. Table S5 for the full list of significant GO terms.

**Figure S10.**
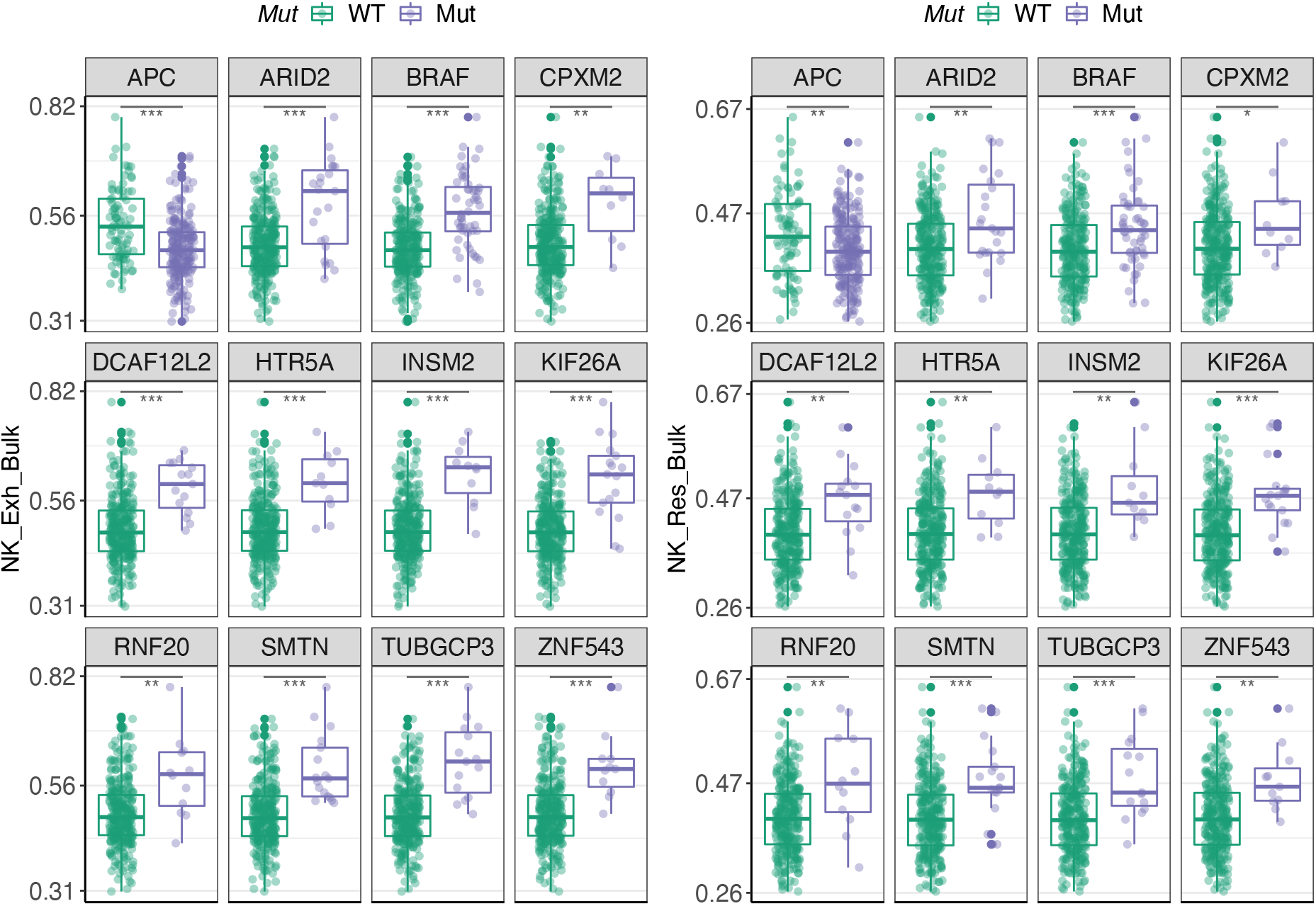
Elastic net results – genes whose mutations are predictive of both NK_Exh_Bulk and NK_Res_Bulk.

**Figure S11.**
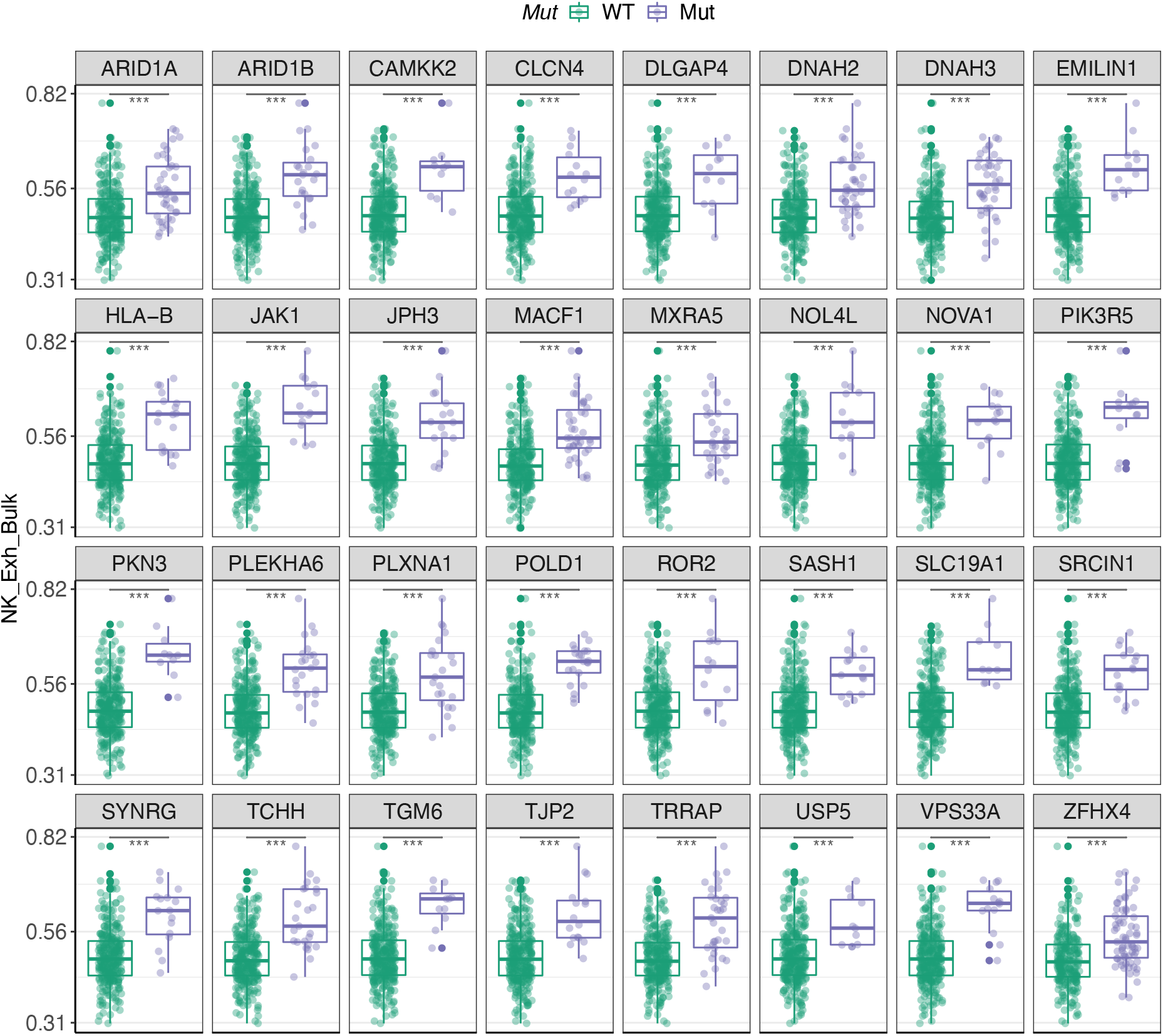
Elastic net results – genes whose mutations are predictive of NK_Exh_Bulk.

**Figure S12.**
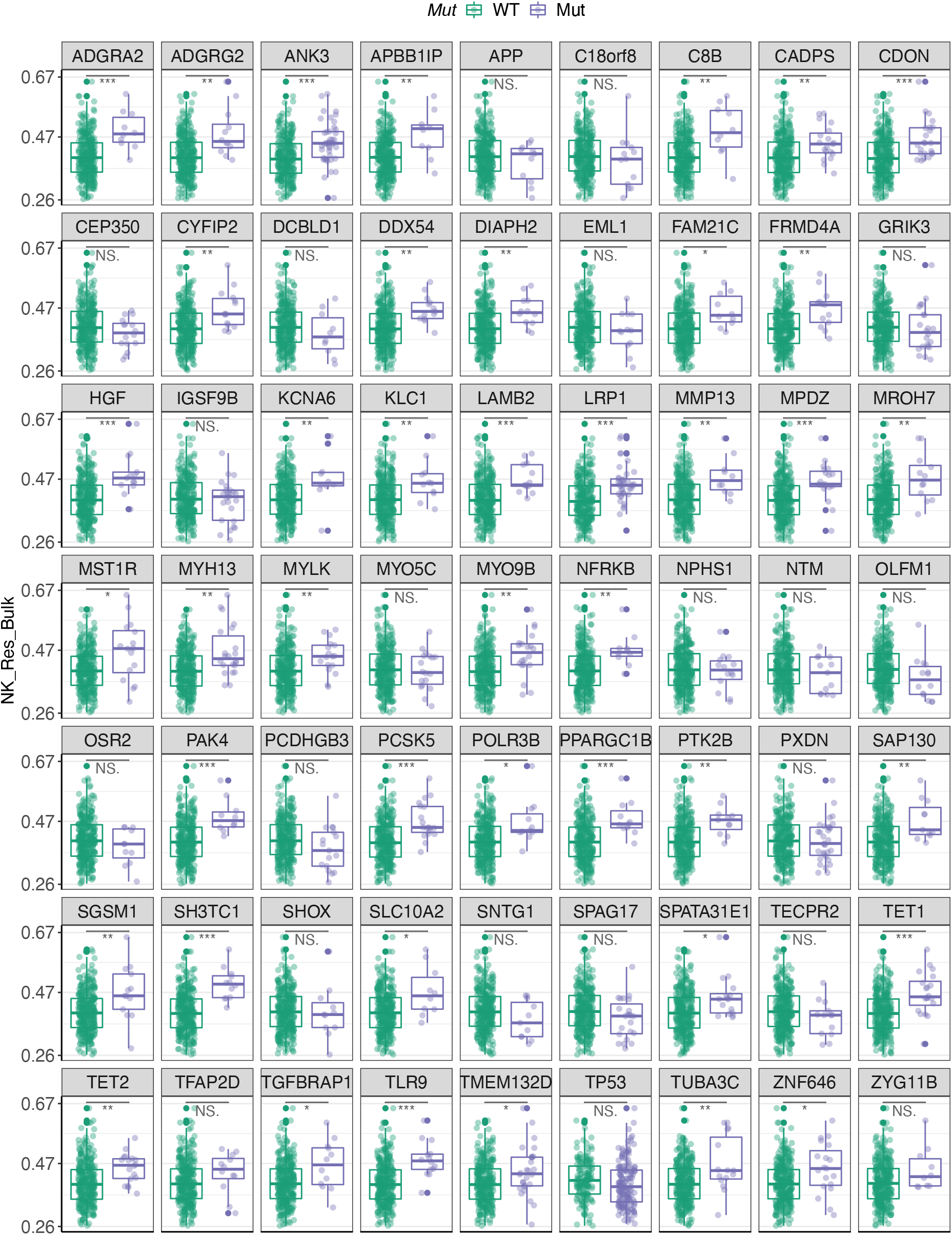
Elastic net results – genes whose mutations are predictive of NK_Res_Bulk.

**Figure S13.**
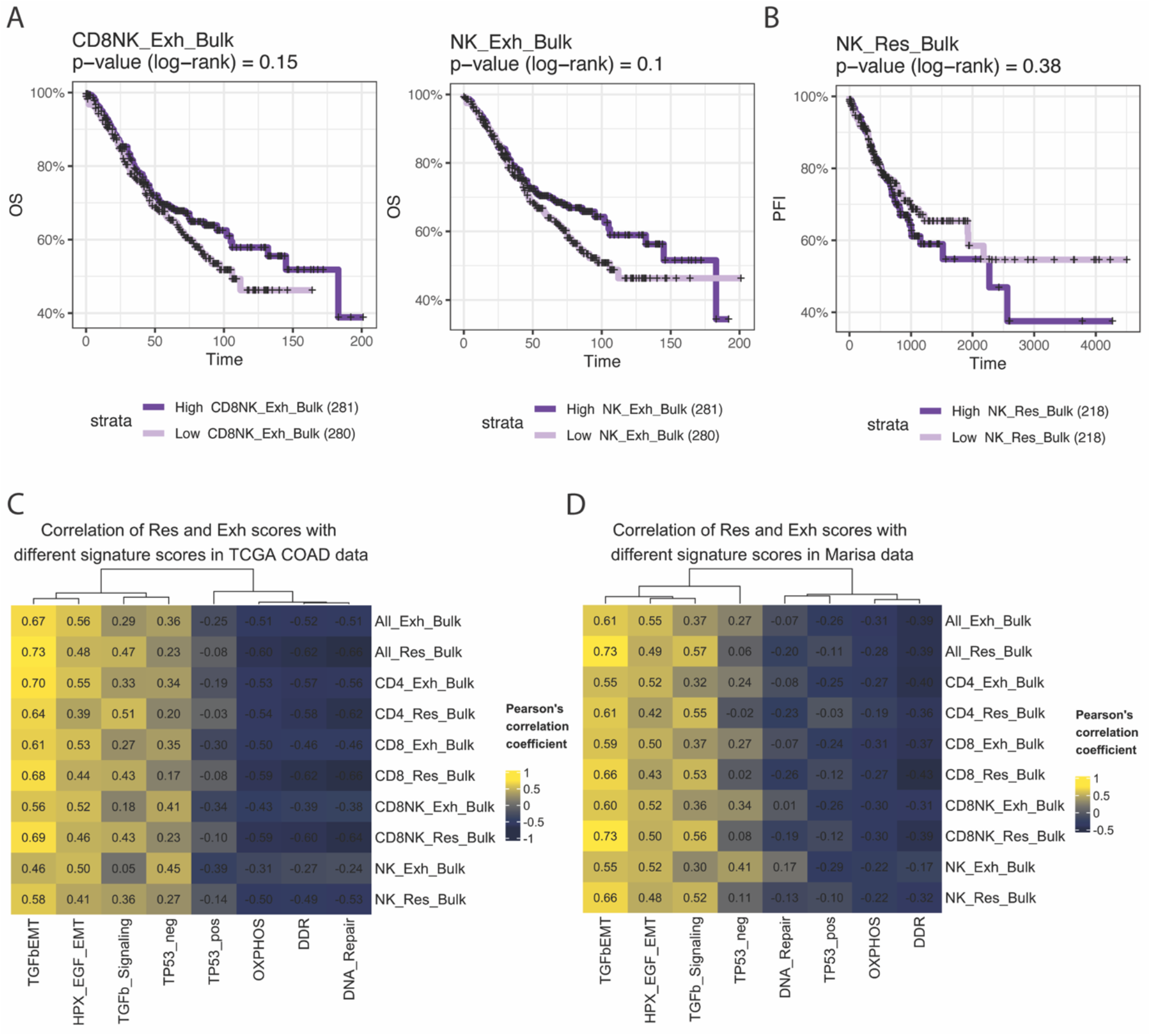
**A.** Kaplan-Meier curves demonstrating overall survival (OS) for patients from Marisa data stratified to samples with high and low scores based on median score values (multivariate Cox proportional hazard model *p*-value < 0.05 for both signatures). **B.** Kaplan-Meier curves demonstrating progression free interval (PFI) for patients from TCGA data stratified to samples with high and low scores based on median score value (multivariate Cox proportional hazard model *p*-value < 0.05). **C & D.** Pearson’s correlation coefficients between Res_Bulk or Exh_Bulk and several cancer related signatures in TCGA (**C**) and Marisa (**D**) data.

**Figure S14.**
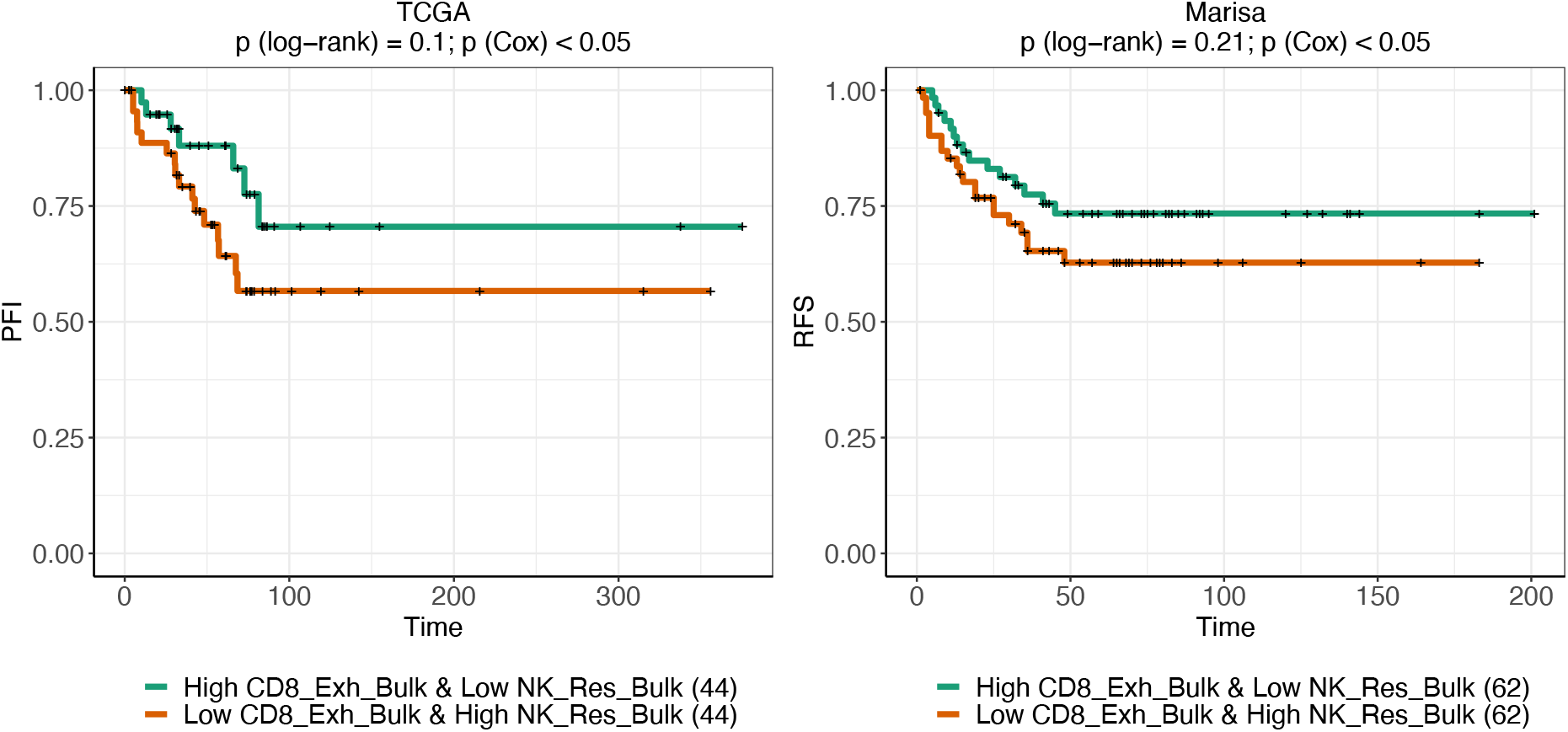
PFI and RFS for samples with high CD8_Exh_Bulk & low NK_Res_Bulk vs those with low CD8_Exh_Bulk & high NK_Res_Bulk. Although the log-rank tests were not significant, the multivariate Cox regression showed significant results in both datasets.

