## Supplemental Table S7 for "The ratio of exhausted to resident infiltrating lymphocytes is prognostic for colorectal cancer patient outcome"

**Supp. Table S7.** List of the tools and packages used in the manuscript: Foroutan et al “The ratio of exhausted to resident infiltrating lymphocytes is prognostic for colorectal cancer patient outcome”.

| Tool/package | Version | Refernce |
| --- | --- | --- |
| Seurat | 3.2.2 | [1] |
| singscore | 1.8.0 | [2] |
| SingleCellExperiment | 1.10.1 | [3] |
| slingshot | 1.6.1 | [4] |
| sctransform | 0.3 | [5] |
| PMA | 1.2.1 | https://cran.r-project.org/web/packages/ PMA |
| MAST | 1.14.0 | Bioconductor package; DOI: 10.18129/B9.bioc.MAST |
| GEOquery | 2.56.0 | [6] |
| RUV-III (ruv package) | 0.9.7.1 | [7], https://cran.r-project.org/web/packages/ruv |
| CMScaller | 0.99.2 | [8] |
| msigdf | 5.2 | https://github.com/stephenturner/msigdf |
| TCGAbiolinks | 2.16.1 | [9] |
| tidyverse | 1.3.0 | [10] |
| edgeR | 3.30.3 | [11] |
| limma | 3.44.3 | [12] |
| survival | 3.2-3 | https://cran.r-project.org/web/packages/survival |
| survminer | 0.4.8 | https://cran.r-project.org/web/packages/survminer |
| ggfortify | 0.4.10 | https://cran.r-project.org/web/packages/ggfortify |
| foreach | 1.5.0 | https://cran.r-project.org/web/packages/foreach |
| ggsignif | 0.6.0 | https://cran.r-project.org/web/packages/ggsignif |
| RColorBrewer | 1.1-2 | https://cran.r-project.org/web/packages/RColorBrewer |
| ComplexHeatmap | 2.4.3 | [13] |
| broom | 0.7.0 | [14] |
| glmnet | 4.0-2 | [15] |
| gridExtra | 2.3 | https://cran.r-project.org/web/packages/gridExtra |
| GO.db | 3.11.4 | Bioconductor package; DOI: 10.18129/B9.bioc.GO.db |
| rrvgo | 1.1.2 | Bioconductor package; DOI: 10.18129/B9.bioc.rrvgo |
| org.Hs.eg.db | 3.11.4 | Bioconductor package; DOI: 10.18129/B9.bioc.org.Hs.eg.db |
| hgu133plus2.db | 3.2.3 | Bioconductor package; DOI: 10.18129/B9.bioc.hgu133plus2cdf |
| annotate | 1.66.0 | Bioconductor package; DOI: 10.18129/B9.bioc.annotate |
| SummarizedExperiment | 1.18.2 | Bioconductor package; DOI: 10.18129/B9.bioc.SummarizedExperiment |
