## Supplemental Tables S8 and S9 for "The ratio of exhausted to resident infiltrating lymphocytes is prognostic for colorectal cancer patient outcome"

**Supp. Tables S8 and S9 associated with the manuscript: Foroutan et al “The ratio of exhausted to resident infiltrating lymphocytes is prognostic for colorectal cancer patient outcome”.**

**Supp. Table S8.** Studies identifying genes associated with Exh/Res programs

| Used in cell type | Signature name | Context | Species | Reference |
| --- | --- | --- | --- | --- |
| T and NK | CD8_TEX_Guo | NSCLC | Human | [1] |
| T and NK | CD4_TEX_Guo | NSCLC | Human | [1] |
| T and NK | CD8_TEX_JA | Melanoma | Human | [2] |
| T and NK | CD4_TEX_JA | Melanoma | Human | [2] |
| T and NK | CD8_Dysfunc_Li | Melanoma | Human | [3] |
| T and NK | CD4_TregsTum_Li | Melanoma | Human | [3] |
| T and NK | TRM_review | Cancer | Human and mouse | [4] |
| T and NK | TEX_ImmuneSig | Cancer | Human | Manual curation |
| T and NK | TerminallyDiff_T | Cancer | Human | [5] |
| T and NK | CorridoniGenes; subset to markers of Trm, IL26, Double Positive (DP), IEL (TYROBP+/-), and GZMK-eff (2) | Colitis | Human | [6] |
| T and NK | lungSpleenTRM | Normal | Human | [7] |
| T and NK | Tex, Tpex, ExhCore, Trm | Acute and chronic viral infections | Mouse | [8] |
| T and NK | sigsTRM_tissue (for gut, lung, skin) | HSV and LCMV infected tissues | Mouse | [9] |
| T and NK | sigsTRM_tissue (liver) | irradiated sporozoites | Mouse | [10] |
| T and NK | sigsTRM_tissue (brain) | Acute virus infection | Mouse | [11] |
| NK | CD65bright, CD56dim, CD56dimCD57pos, CD56dimCD57neg | Normal tonsil and blood | Human | [12] |
| NK | ILC1 | Normal tonsil and intra-epithelial cells | Human | [12] |
| NK | ILC1 | Cancer | Mouse | [13] |
| NK | NK_vs_ILCs | Normal tonsil | Human | [14] |
| NK | ILC1 (ILC1_vs_ILC2.3) | Normal tonsil | Human | [14] |
| NK | CD69pCD49apCD103p_vs_CD69n,  CD69pCD49apCD103n_vs_CD69n,  CD69sp_vs_CD69n,  CD69pCD49apCD103p_vs_CD69sp,  CD69pCD49apCD103n_vs_CD69sp | Normal lung, spleen, bone marrow | Human | [7] |
| NK | resident3Genes,  residentGenes,  residentNK_lungBM_notTRM,  residentNK_lungOnly | Normal lung, spleen, bone marrow | Human | [7] |
| NK | NKGenes (from a review paper) | Several normal tissues | Human | [15] |
| NK | NK_ImmGene,  NK_cur,  NK_Xiong,  NK_imsig,  NK_JA | Several general NK signatures | Human | [2, 16-18] |

**Suppl Table S9.** List of signatures used for scoring tumour data and performing survival analysis, as well as signatures used to score single cell data. The HALLMARK signatures were obtained from MSigDB [19] through msigdf R package.

| Signature | ref |
| --- | --- |
| HALLMARK_REACTIVE_OXIGEN_SPECIES_PATHWAY | msigdf R package |
| HALLMARK_OXIDATIVE_PHOSPHORYLATION | msigdf R package |
| HALLMARK_PEROXISOME | msigdf R package |
| HALLMARK_KRAS_SIGNALING_UP | msigdf R package |
| HALLMARK_P53_PATHWAY | msigdf R package |
| HALLMARK_WNT_BETA_CATENIN_SIGNALING | msigdf R package |
| HALLMARK_TGF_BETA_SIGNALING | msigdf R package |
| HALLMARK_GLYCOLYSIS | msigdf R package |
| HALLMARK_HYPOXIA | msigdf R package |
| HALLMARK_DNA_REPAIR | msigdf R package |
| PID_NFAT_TFPATHWAY | msigdf R package |
| HALLMARK_INTERFERON_ALPHA_RESPONSE | msigdf R package |
| HALLMARK_INTERFERON_GAMMA_RESPONSE | msigdf R package |
| HALLMARK_IL6_JAK_STAT3_SIGNALING | msigdf R package |
| HALLMARK_EPITHELIAL_MESENCHYMAL_TRANSITION | msigdf R package |
| TP53-negative and TP53-positive | [20] |
| HPX_EGF_EMT (Hypoxia/EGF-induced EMT) | [21] |
| TGFbEMT (TGF-β induced EMT) | [22] |
| Epi_Thiery and Mes_Thiery | [23] |
| human DNA damage repair (DDR) | [24] |
| Hypoxia_Buffa | [25] |
| MYC-negative and MYC-positive | [26] |
| TRM_TGFb_Unstim_Up (TGF-β signature in T_rm_ cells) | [27] |
| Proliferation_ImSig | [17] |
| Antigen presentation (IFNGR1, IFNGR2, TAP1, TAP2, TAPBP, JAK1, JAK2, STAT1, CASP8, and B2M) | Manual curation |
